# *Legionella pneumophila* infection and antibiotic treatment engenders a highly disturbed pulmonary microbiome with decreased microbial diversity

**DOI:** 10.1101/808238

**Authors:** Ana Elena Pérez-Cobas, Christophe Ginevra, Christophe Rusniok, Sophie Jarraud, Carmen Buchrieser

**Author notes:** For correspondence: Carmen Buchrieser, Institut Pasteur, Biologie des Bactéries Intracellulaires, 28, rue du Dr. Roux, 75724 Paris Cedex 15, France.

## Abstract

**Background:** Lung microbiome analyses have shown that the healthy lung is not sterile but it is colonized like other body sites by bacteria, fungi and viruses. However, little is known about the microbial composition of the lung microbiome during infectious diseases such as pneumonia and how it evolves during antibiotic therapy. To better understand the impact of the composition of the pulmonary microbiome on severity and outcome of pneumonia we analysed the composition and evolution of the human lung microbiome during pneumonia caused by the bacterium *Legionella pneumophila*.

**Results:** We collected 10 bronchoalveolar lavage (BAL) samples from three patients during long-term hospitalisation due to severe pneumonia and performed a longitudinal in-depth study of the composition of their lung microbiome by high-throughput Illumina sequencing of the 16S rRNA gene (bacteria and archaea), ITS region (fungi) and 18S rRNA gene (eukaryotes). We found that the composition of the bacterial lung microbiome during pneumonia is hugely disturbed containing a very high percentage of the pathogen, a very low bacterial diversity, and an increased presence of opportunistic microorganisms such as species belonging to Staphylococcaceae and Streptococcaceae. The microbiome of antibiotic treated patients cured from pneumonia represented a different perturbation state with a higher abundance of resistant bacteria (mainly Firmicutes) and a significantly different bacterial composition as that found in healthy individuals. In contrast, the mycobiome remains more stable during pneumonia and antimicrobial therapy. Interestingly we identified possible cooperation within and between both communities. Furthermore, archaea (*Methanobrevibacter*) and protozoa (*Acanthamoeba* and *Trichomonas*) were detected.

**Conclusions:** Bacterial pneumonia leads to a collapse of the healthy microbiome and a strongly disturbed bacterial composition of the pulmonary microbiome that is dominated by the pathogen. Antibiotic treatment allows some bacteria to regrow or recolonize the lungs but the restoration of a healthy lung microbiome composition is only regained a certain time after the antibiotic treatment. Archaea and protozoa should also be considered, as they might be important but yet overseen members of the lung microbiome. Interactions between the micro- and the mycobiome might play a role in the restoration of the microbiome and the clinical evolution of the disease.

## BACKGROUND

Historically the lower airways were thought to be sterile unless infected, however, the development of culture-independent techniques and high-throughput sequencing have revealed that the respiratory tract harbours like almost every mucosal surface in the human body a distinct microbiome composed of bacteria, fungi and viruses [1–3]. In 2016, the bacterial composition of the healthy lung was the first time deeply characterized by analysing bronchoalveolar lavage fluid (BAL) samples from healthy individuals. Two types of microbiome profiles (termed pneumotypes) were described [4]. One of the pneumotypes named supraglottic predominant taxa (SPT) was characterized by a high bacterial load and the presence of anaerobes such as *Prevotella* and *Veillonella* belonging to the phyla Firmicutes and Bacteroidetes whereas the second pneumotype was called the background predominant taxa (BPT) as it presented a low biomass and identified bacteria belonged mainly to the phyla Proteobacteria, and Actinobacteria such as *Acidocella* or *Pseudomonas*. Although biology of the lung microbiome is a new field and very little is known about its role in human health, it was suggested that the lung microbiome participates in immune system functions including immune cell maturation and inflammatory responses [2, 5]. Segal and colleagues proposed that bacteria such as *Prevotella* and *Veillonella* that have been commonly identified in the lungs of healthy individuals or their products participate in the regulation of inflammation and are linked to the activation of the lung mucosal Th17 response [4].

The healthy lung mycobiome is mainly composed of environmental fungi belonging to the phyla Ascomycota and Basidiomycota such as Davidiellaceae, *Eurotium Eremothecium*, and *Cladosporium*, while during disease a higher abundance of pathogenic species belonging to the genera *Aspergillus*, *Malassezia* or *Candida* are present [6]. Commensal fungi, like bacteria, seem to participate in immune system stimulation, inflammatory response and protection against pathogens [5, 6]. In contrast to the lung micro- and mycobiome that starts to be investigated better, the presence of eukaryotes, archaea and viruses among the lung microbiota has been very little investigated like for most other body habitats.

To date, the majority of studies analysing the lung microbiome in disease have focused on cystic fibrosis [7–9] asthma [10, 11] or chronic obstructive pulmonary disease (COPD) [12–14]. These studies have shown that lung microbial dysbiosis is generally characterized by a microbiome enriched in pathogenic and opportunistic bacteria (such as certain Gammaproteobacteria), with a high biomass due to the overgrowth of pathogens and a low diversity because of the displacement of the normal members of the community [2, 5]. However, few studies have addressed the role of the microbiome composition during infectious diseases in the lungs.

Acute lower respiratory tract infections are the leading cause of morbidity and mortality worldwide among infectious diseases. Among these, pneumonia represents a clinical and economic burden and a major public health problem, particularly for children and the elderly. *Legionella pneumophila* is one of these human, bacterial pathogens as it is causing a severe pneumonia called Legionnaires’ disease that is often fatal when not treated promptly. It is characterized by clinical polymorphism and variable severity, 98% of cases are hospitalized and 40% require intensive care unit (ICU) admission. The mortality rate remains high (11% to 33% in ICUs) despite improved diagnostic and therapeutic management of patients. The characterization of the pulmonary microbiome during *Legionella* infection has been poorly addressed. Recently, Mizrahi and collaborators characterized the microbiome of nine sputum samples from patients with *L. pneumophila* infection finding a higher presence of *Streptococcus* and a low presence of *L. pneumophila* [15], but lung samples like pulmonary lavage or tracheal aspirates have not been analysed. Furthermore, no study has analysed the evolution of the pulmonary microbiome and mycobiome during *L. pneumophila* infection and antibiotic treatment although antibiotics are one of the most important factors in reshaping the microbiome composition with important consequences for health [3].

Here we report a comprehensive analysis of the lung microbiome composition of three patients with long-term pneumonia due to *L. pneumophila* and its evolution during antibiotic treatment. Using high-throughput sequencing of marker genes we show that the pulmonary microbiome is highly disturbed during pneumonia and by the antibiotic treatment whereas the mycobiome is less affected. Our analyses included also archaea and eukaryotes (amoeba) showing that both are present in the pulmonary microbiota and that they might also play a role in the response to the microbiome disturbance.

## RESULTS AND DISCUSSION

### The analysed human lung samples originated from patients and healthy individuals

Patient A is a 28-year-old immunocompetent man that was admitted to the hospital due to severe community-acquired pneumonia (CAP). The first five days of hospitalisation the patient was treated with amoxicillin as a *Streptococcus pneumoniae* infection was suspected. As the pneumonia got worse this treatment was enlarged with amoxicillin/clavulanic and spiramycin and later with ceftriaxone, erythromycin, levofloxacin and Tamiflu for severe CAP. Only at day five *L. pneumophila* was identified as the causative agent and the antibiotic treatment was switched to a combination of erythromycin and levofloxacin [16, 17] (**Figure 1A**). During one month and a half of hospitalization and treatment the patient was sampled intensively and due to the suspicion of the presence of a lung abscess according a thoracic computed tomography scan (CT-SCAN), rifampicin was added at day 13. While the patient remained febrile, a second thoracic CT-SAN indeed revealed a voluminous lung abscess at day 34 and *Fusobacterium nucleatum* was identified in addition to *L. pneumophila.* Thus, nitroimidazole was added for 20 more days. The abscess was resected at day 42 of treatment and the patient recovered fully. Here we analysed the microbiome composition of BAL samples taken at day 5, 14, 24, 34 and 42 post-admission as well as one sputum sample taken at day 4 post-admission and a biopsy taken during the lung abscess removal at day 42 (**Figure 1A**).

**Figure 1.**
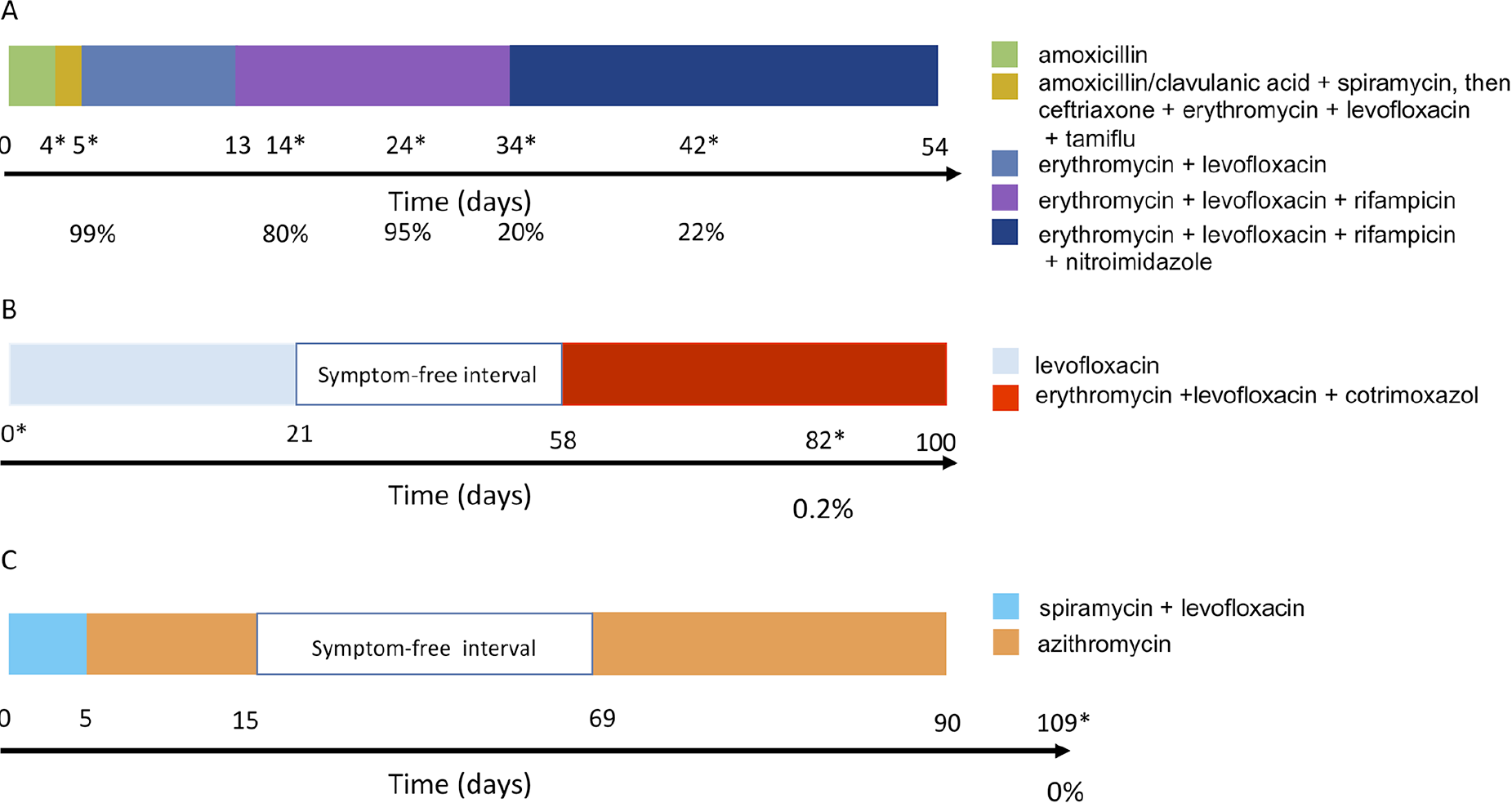
History of the antibiotic treatment of the patients analysed here. (A) Patient A (B) patient B, (C) patient C. The colours indicate the antibiotics treatment and the length of the coloured block the duration of the treatment with each antibiotic. The numbers below the coloured blocks indicate the days where a sample was taken. The BAL samples for which the microbiome was analysed here are marked with a star. The percentage given below the time line indicates the relative abundance of *Legionella* present in the sample as determined by qPCR.

Patient B, a 69-year-old immunocompromised woman, was hospitalised for pneumonia caused by *Legionella* and treated for 21 days with levofloxacin. After a period of 37 days free of symptoms, she was re-hospitalised for bilateral pulmonary consolidations and pleural effusion. A recurrent *Legionella* caused pneumonia was confirmed and she was treated for six weeks with a combination of erythromycin and levofloxacin and also cotrimoxazole for a concomitant pneumocystosis [17]. A sample was taken at the onset of the treatment (day 0) and after two months and a half (day 82) (**Figure 1B**). We analysed both samples to study the composition of the microbial community composition at the beginning and the end of the treatment.

Patient C, a 76-year-old immunocompromised man, was sampled 109 days after the onset and 19 days after the end of the therapy when the patient was fully recovered; this BAL was negative for *L. pneumophila* as confirmed by PCR [17]. Thus, this sample represents the lung microbiome after an intensive antibiotic treatment (named C109) (**Figure 1C**).

To compare the pneumonia microbiomes with that of healthy lung microbiomes we retrieved the raw data of a recent study published by Segal and colleagues where the lung microbiome of 49 healthy individuals was characterized by sequencing the V4 region of the 16S rRNA gene obtained from BAL samples [4] (accession number Gene Expression Omnibus (GEO): GSE74395).

### During pneumonia the pulmonary microbiome is largely dominated by *Legionella*

The bacterial composition of the BAL samples of patient A was clearly dominated by *Legionella* during the first 24 days of the antibiotic treatment (**Figure 2A**). At the time of diagnosis *Legionella* represented 99% of the identified bacteria in the lungs. After 10 days of adapted treatment (day 14) *Legionella* was still responsible for around 80% of the bacterial composition, followed by *Enterococcus* with a relative abundance of 13%. Interestingly, at day 24 *Legionella* increased again to a relative abundance of 95% of the total amount of bacteria. Similarly, the microbiome of patient B was mainly composed of *Legionella* representing 57% of the diversity at the beginning of the treatment (**Figure 2B**). Thus, *Legionella* represents the vast majority (57-99%) of the bacterial community during infection similar to what was shown for *Pseudomonas aeruginosa* infections [18, 19].

**Figure 2.**
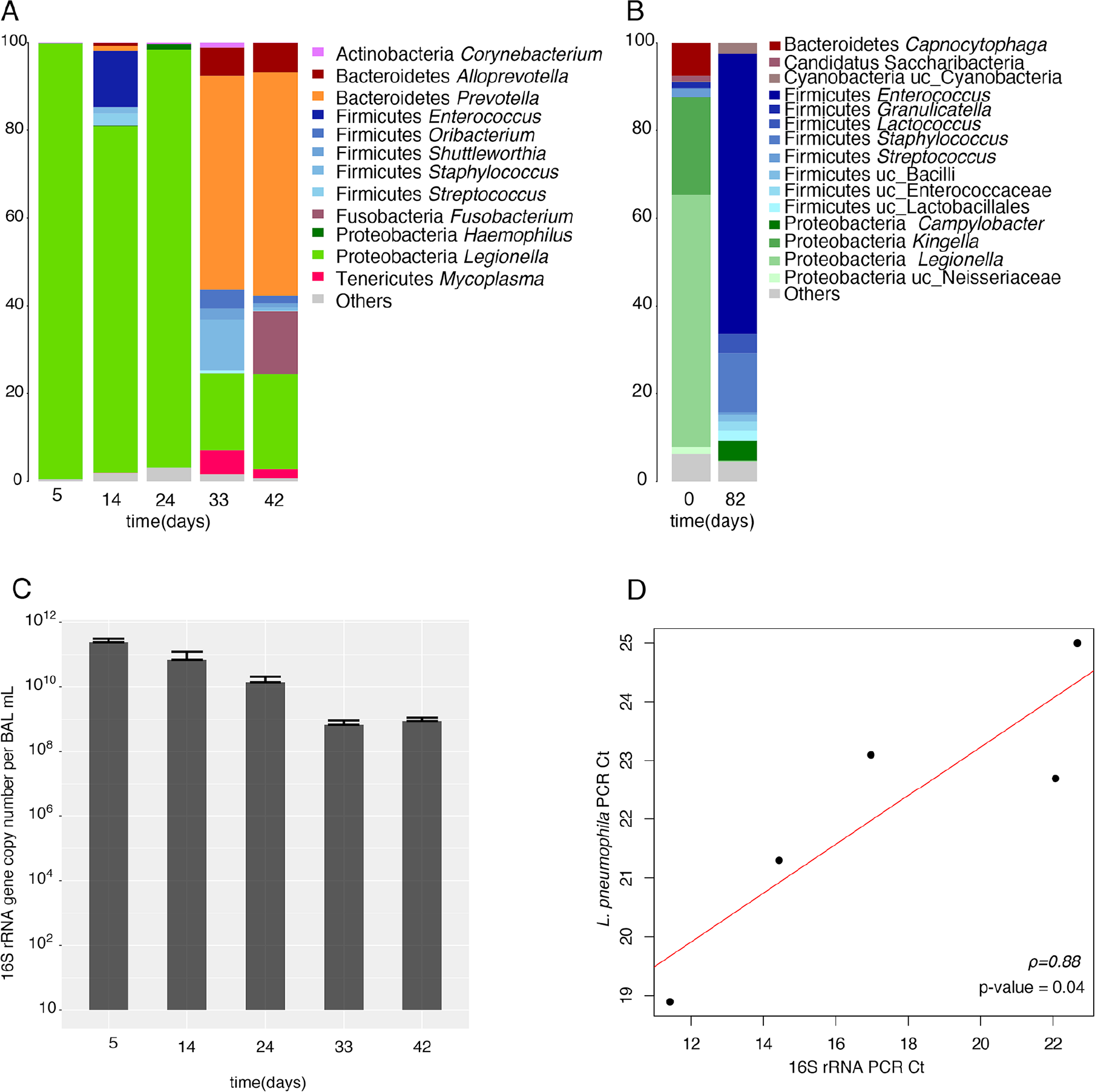
Microbiome composition of the BAL samples. **(A)** Bacterial composition of the BAL samples of patient A. The taxonomy is based on the RDP. **(B)** Bacterial composition of the BAL samples of patient B. The taxonomy is based on the RDP. **(C)** Number of total bacteria (patient A) estimated by qPCR of the 16S rRNA gene. **(D)** Correlation of the number of bacteria (Ct of the 16S rRNA gene qPCR) with the amount of *Legionella* (Ct of the *mip* gene qPCR) (patient A). The Ct values for the *mip* gene were obtained from a clinical case-study [16].

### Antibiotic treatment leads to strong perturbations and slow recovery of a healthy microbiome

A strong change occurred in the microbiome composition of patient A between day 24 and 34 probably due to the changed treatment (30 days after the beginning of treatment and 20 days after adding rifampicin) (**Figure 2A**). First, we found a marked decrease in the relative abundance of *Legionella*, representing only around 20% of the composition. Secondly, we found a recovery of the relative abundance of certain bacteria commonly found in the lung microbiome of healthy individuals. Importantly, the abundance of *Prevotella* changed from less than 1% during the first 24 days of therapy to around 50% at day 34. Indeed, certain species of the genus *Prevotella* are resistant to different classes of antibiotics including macrolides, which were included in the treatment [20]. Furthermore, *Prevotella* was described as predominant taxa in healthy microbiomes [4] suggesting that the microbiome started to recover. At day 33 additional normal bacteria of the lung microbiome such as *Staphylococcus* (11%), *Alloprevotella* (6%) and *Oribacterium* (4%) were identified. Finally, an increase in the abundance of *Fusobacterium* (14%) was observed. Interestingly, *F. nucleatum* is ubiquitous in the oral cavity, but absent or rarely detected at other body sites of healthy people. However, under disease conditions, this bacterium is one of the most prevalent species found in extra-oral sites [21]. Erythromycin (macrolide) is also not effective against *Fusobacterium* explaining the increase of its abundance in the last days of treatment [22].

The species richness of the different samples was estimated based on the number of observed OTUs and the richness estimator Chao 1, while the species evenness was based on the Shannon index. According to these metrics, the bacterial diversity collapsed due to the antibiotic treatment around day 24 reaching the lowest values for all estimated parameters (**Supplementary figure 1A**). A collapse in bacterial diversity during antibiotic treatment has also been described for other human microbial communities such as the gut [23, 24].

The bacterial load analysis based on qPCR of the 16S rRNA gene showed a decrease in biomass during the antibiotic treatment (**Figure 2C**). During the first days of treatment the bacterial load was very high (10^10^ - 10^12^ copies of the 16S rRNA gene) but after day 24 the load decreased to 10^8^-10^9^ probably due to the addition of rifampicin at day 13 and/or to the time needed for activity of the combination of fluoroquinolone plus macrolide. The difference between the days 5 and 14 compared to days 34 and 42 was statistically significant (Wilcoxon signed-rank test p-value=0.002). Our Pearson correlation analysis between the Ct values of *L. pneumophila* and the total biomass (16S rRNA data) pointed to a significant positive correlation between both (p-value = 0.04) (**Figure 2D**). Thus, the decrease in biomass can be partially explained by the decrease of the pathogen abundance due to the antibiotic therapy.

The lung microbiome of patient B showed a similar dynamics. At the beginning of the long-term treatment (day 0) with several levofloxacin, erythromycin and cotrimoxazole, (Figure 1B), the microbiome was mainly composed of *Legionella* (57%) and secondly of *Kingella* (22%) and *Capnocytophaga* (7.6%) (**Figure 2B**), genera that are not reported to be part of bacteria present in the healthy lungs [4].. After two long periods of antibiotic treatment (day 82) the composition changed dramatically, and the most abundant bacteria then were *Enterococcus* (64%), *Staphylococcus* (14%), *Campylobacter* (4.5%) and *Lactococcus* (4.4%), while the abundance of *Legionella* decreased to 0.2% and *Kingella* and *Capnocytophaga* were not present anymore. The enrichment of *Enterococcus* after the treatment might be due to the often-observed resistance of these bacteria to several different antibiotics including fluoroquinolones, macrolides and cotrimoxazole, and as described for the gut [25], the antibiotic treatment may also lead to the dissemination of enterococci in the lungs. Furthermore, after antibiotic treatment we observed, like for patient A, an increase in *Staphylococcus*, a genus reported to be frequently resistant to fluoroquinolones and macrolides [26]. The species evenness based on the Shannon index was higher during infection than after treatment, but like for patient A, the species richness was higher at the end of the infection (**Supplementary Figure 1B**).

### During bacterial pneumonia and antibiotic therapy the mycobiome is less disturbed than the microbiome

The mycobiome composition during long-term legionellosis and antibiotic treatment (Patient A) showed different dynamics than the microbial composition. The fungal microbiome fraction was mainly composed of species belonging to the phyla Basidiomycota, and Ascomycota (**Figure 3A**). These two phyla are reported to be the most abundant ones in the human respiratory tract [6]. The major taxa that we identified during the entire sampling period were unclassified species belonging to Agaricomycetes (16-50%), Ascomycota (20-44%), Agaricales (10-18%) and Basidiomycota (8-16%). This indicates that a high diversity of fungi from the human mycobiome remains unknown and awaits characterisation. Some of the most abundant genera identified were *Candida* (0.8-7%) and *Morchella* (0.4-3.5%) (Ascomycota) and *Malassezia* (0.2-4%) (Basidyomycota). Despite the higher homogeneity of the mycobiome composition, the richness and diversity of the fungal species showed a parallel dynamics as it decreased also through the antibiotic treatment, reaching the minimum at day 24th of treatment similar to the microbiome (**Supplementary figure 2A**). Again, as seen for the bacterial fraction the fungal diversity started to recover by day 34 of the treatment. The changes in the mycobiome during bacterial infection and antibiotic treatment suggest that the fungal and the bacterial communities interact ecologically in the lungs (i.e. cooperation, competition) and with the host immune system and the inflammatory responses. Indeed, in a mouse model of oropharyngeal candidiasis a potential mutualism has been reported where the main pathogen *Candida albicans* promotes mucosal biofilm formation and colonization of *Streptococcus oralis* through modulation of the mouse inflammatory response, thus increasing the virulence of the infection [27]. Also, *C. albicans* favours the growth of *Pseudomonas aeruginosa* during pneumonia by affecting the production of reactive oxygen species (ROS) by alveolar macrophages in rats [28]. Thus, also in the human lung the displacement of some key microorganisms in the microbial community due to infection and treatment may affect the survival of other microorganisms that depend on them.

**Figure 3.**
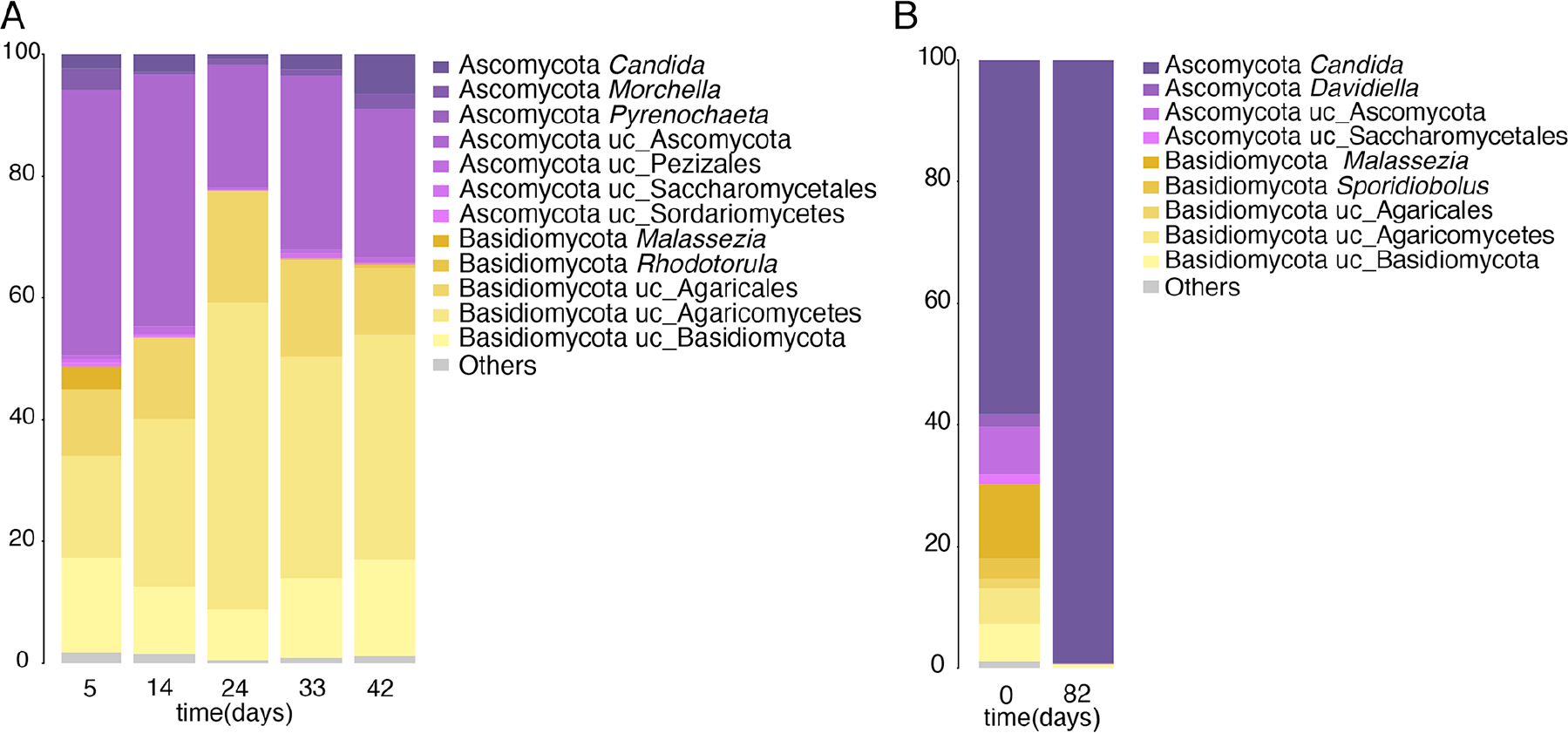
Mycobiome composition of the BAL samples. **(A)** Fungal composition of the BAL samples of patient A. The taxonomy is based on the Warcup ITS training set. The phyla and genera are shown for the most abundant groups (>1%). **(B)** Fungal composition the BAL samples of patient B. The taxonomy is based on the Warcup ITS training set. The phyla and genera are shown for the most abundant groups (>1%).

These results were confirmed by the analyses of the mycobiome of patient B. Before antibiotic treatment the genus *Candida* showed a relative abundance of 65% and unclassified fungi from the class Agaricomycetes and Basidiomycota a relative abundance of 11% and 7%, respectively (**Figure 3B**), but after several weeks of antibiotic treatment (day 82) about 99% of the sequences belonged to the genus *Candida.* In parallel to the marked increase of *Candida*, the diversity values (the number of observed OTUs, Chao 1 and Shannon index) were lower after antibiotic therapy (**Supplementary Figure 2B**). Indeed, increase in the abundance of *Candida* has often been associated to extensive antibiotic usage in hospitals [29], similar to what was seen here. Thus, infection and associated antibiotic treatment changes the healthy lung microbiome considerably, and allows *Candida* species to occupy the niche, which is disturbed by the pathogen and the antibiotics treatment.

### BALs represent the lung microbiome composition with higher accuracy than sputum

We compared the BAL sample taken at day 5 with the sputum sample taken at day 4 to compare the difference in the composition (patient A). The most abundant bacterial genera in the sputum were *Streptococcus* (43%), *Prevotella* (13%) and *Gemella* (13%) while *Legionella* represented only 3% of the total relative abundance (**Figure 4A**). In contrast, the BAL sample showed a lower diversity and OTU richness since more than 99% of the sequences belonged to *Legionella* (**Supplementary figure 1A**). This reveals that during pneumonia the presence of *Legionella* is minimal in the sputum while *Legionella* is abundant in the BAL samples that represent the lung. Similarly, the microbiome composition of nine sputum samples from patients with confirmed legionellosis cases detected a relative abundance of *Legionella* of only 3% [15]. Thus, sputum does not represent the lung microbiome composition.

**Figure 4.**
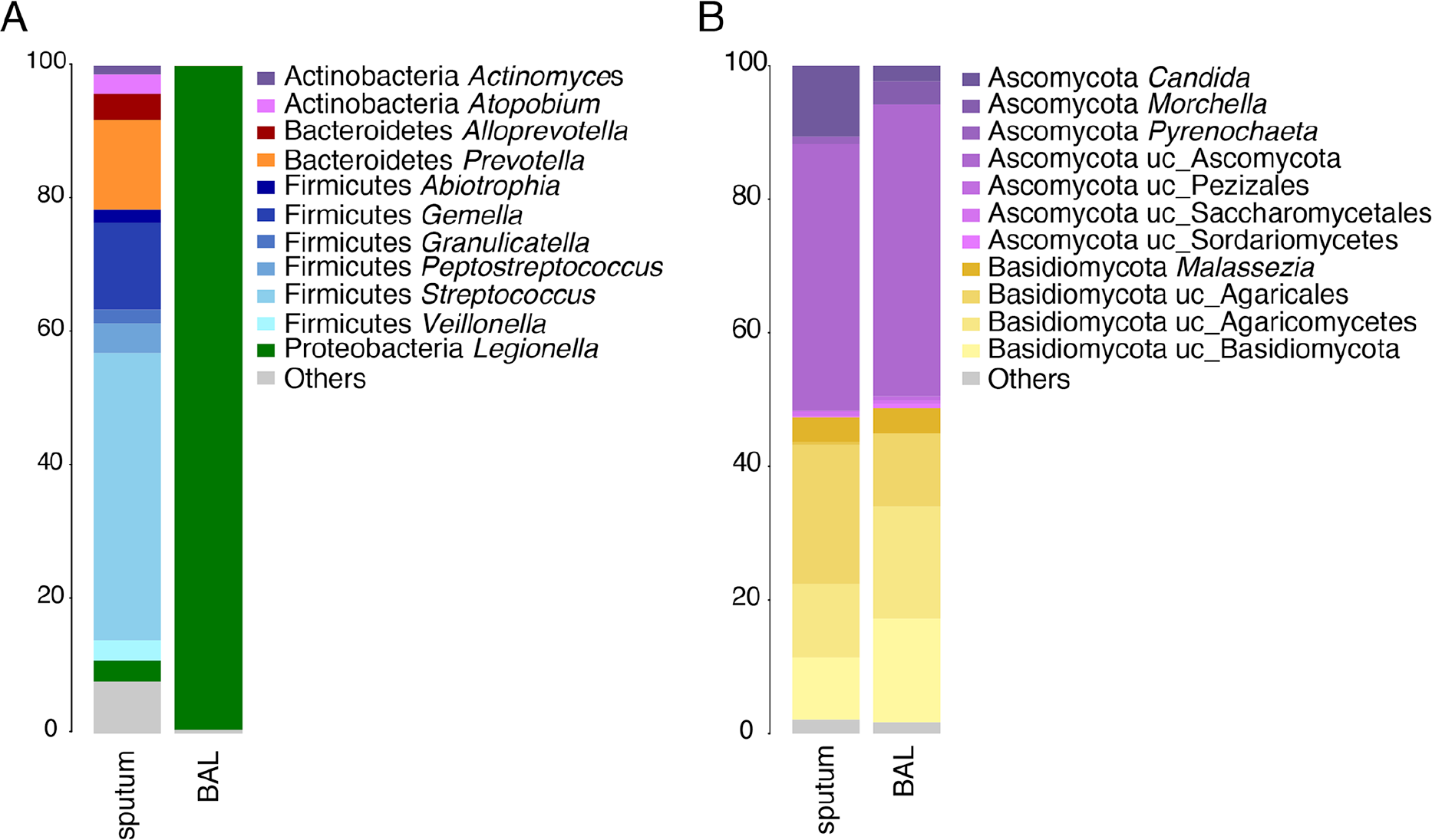
Comparison of the microbiome composition of BAL and sputum samples. **(A)** Bacterial composition. The taxonomy is based on the RDP. **(B)** Fungal composition. The taxonomy is based on the Warcup ITS training set. The phyla and genera are shown for the most abundant groups (>1%).

Interestingly, when the fungal microbiome of the sputum sample and the BAL sample were compared a similar composition of both with a high abundance of Ascomycota (40%), was observed (**Figure 4B**). Slight differences were detected in the abundance of Agaricomycetes (30%), and *Candida* (11%) as these were enriched in the sputum sample. However, the alpha-diversity measures were lower in the sputum than in the BAL sample (**Supplementary figure 2A**) suggesting that bronchoscopy is a more accurate technique for the study of the fungal diversity of the lungs at species level.

### The lung abscess is enriched in pathogenic microorganisms

One of the reasons why patient A did not recover for such a long time despite antibiotic treatment, was probably the presence of a lung abscess that was confirmed only at day 34 and which may have shed *Legionella* constantly into the lungs. We thus characterized also the microbial composition of the lung abscess that was resected by surgery (day 42) (**Figure 5A**). It was mainly composed of *Legionella* (38%), followed by *Prevotella* (19%), *Fusobacterium* (15%) and *Oribacterium* (15%). As expected, *Legionella* was more abundant in the lung abscess than in the BAL samples. In contrast, anaerobic bacteria such as *Fusobacterium, Oribacterium, Shuttleworthia* showed higher abundance in the abscess as the environment is more anoxic [30], but the species richness was lower as less species seemed to be able to colonize the abscess (**Supplementary figure 1A**).

**Figure 5.**
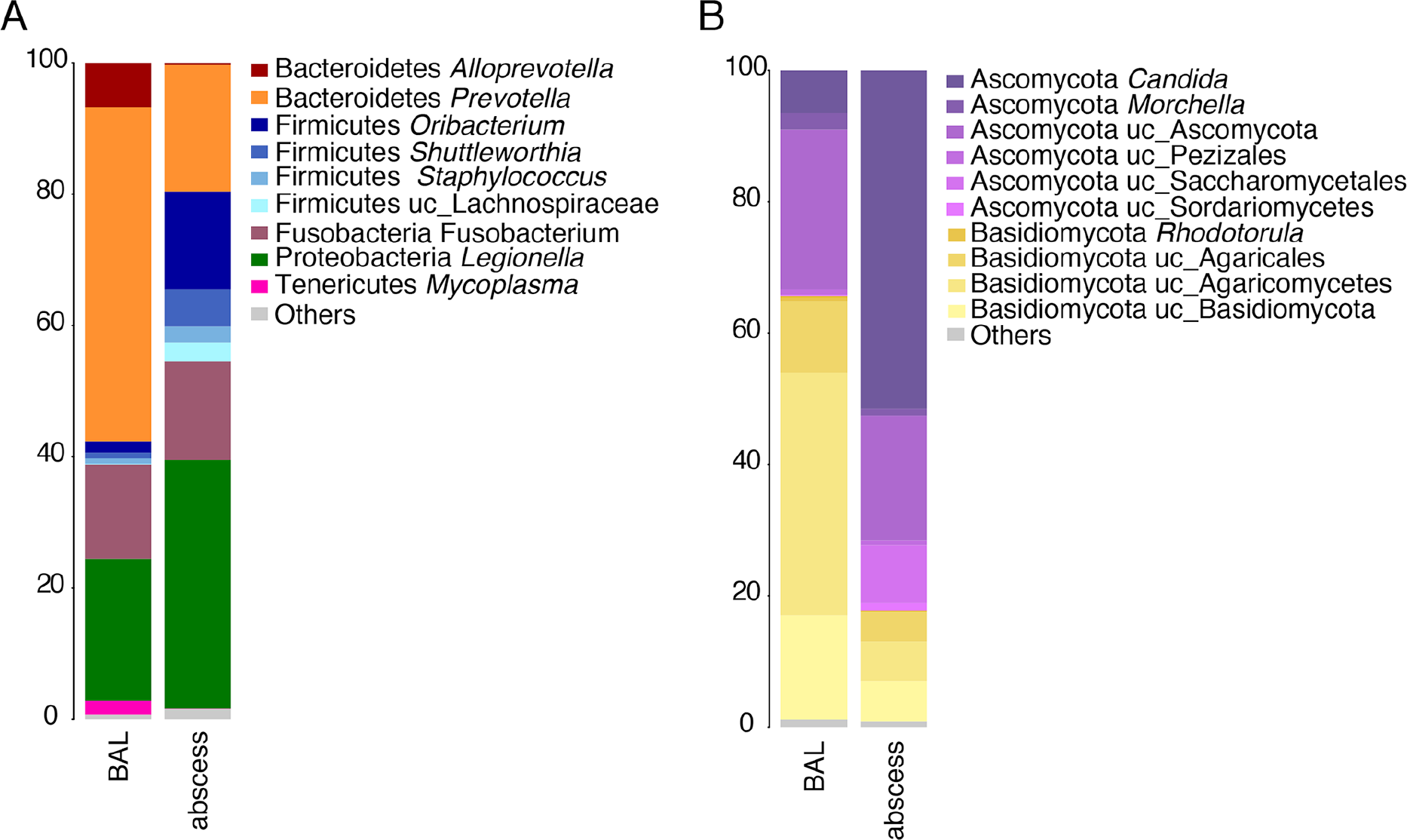
Comparison of the microbiome composition of the BAL and abscess samples. **(A)** Bacterial composition. The taxonomy is based on the RDP. **(B)** Fungal composition. The taxonomy is based on the Warcup ITS training set. The phyla and genera are shown for the most abundant groups (>1%).

The fungal composition of the lung abscess (**Figure 5B**) was dominated by *Candida* (50% of the relative abundance) and other fungal genera belonging to the Ascomycota (20%) revealing enrichment in the relative abundance of this phylum. The species richness was higher in the lung abscess but the Shannon Index was lower as few species were dominant (**Supplementary figure 2A)**. In *in vitro* models it has been shown that *C. albicans* and *Fusobacterium* strongly co-aggregate [31, 32]. Thus, *Candida* may take advantage of the presence of bacterial pathogens such as *F. nucleatus* that was highly abundant in the abscess. Indeed, co-infection in lung abscess of *Candida* and bacteria has been previously reported [33] suggesting that the high abundance of *Candida* is related to the infection with *Legionella* and the presence of *Fusobacterium*.

### After long-term antibiotic treatment the microbiome is enriched in Firmicutes and starts to recover

To analyse the recovery of the microbiome after *L. pneumophila* induced pneumonia and the associated antibiotic treatment we characterized the lung microbiome composition of patient C three weeks after the end of two episodes of antibiotic treatment (spiramycin, levofloxacin, azithromycin) before and after a 55 days symptom-free interval, (**Figure 1C**) defined as day 0 after the end of the treatment.

The bacterial composition was enriched in Firmicutes such as *Streptococcus* (20%), *Prevotella* (17%), *Veillonella* (15%) and *Enterococus* (10%) (**Figure 6A**). The presence of *Prevotella* and *Veillonella* has been identified in the lungs of healthy people suggesting that the lung microbiome of this patient was in the restoration process [4]. The mycobiome was mainly composed of unclassified fungi belonging to the phyla Ascomycota (40%), Basidiomycota (38%) and Agaricomycetes phyla (14%), a fungal composition similar to a healthy microbiome (**Figure 6B**). Similarly, to the samples from patient B the species evenness (Shannon=4.9) and OTU richness (Chao 1=2348, number of OTUs= 990) after treatment were higher, compared to the samples from patient A and B during infection.

**Figure 6.**
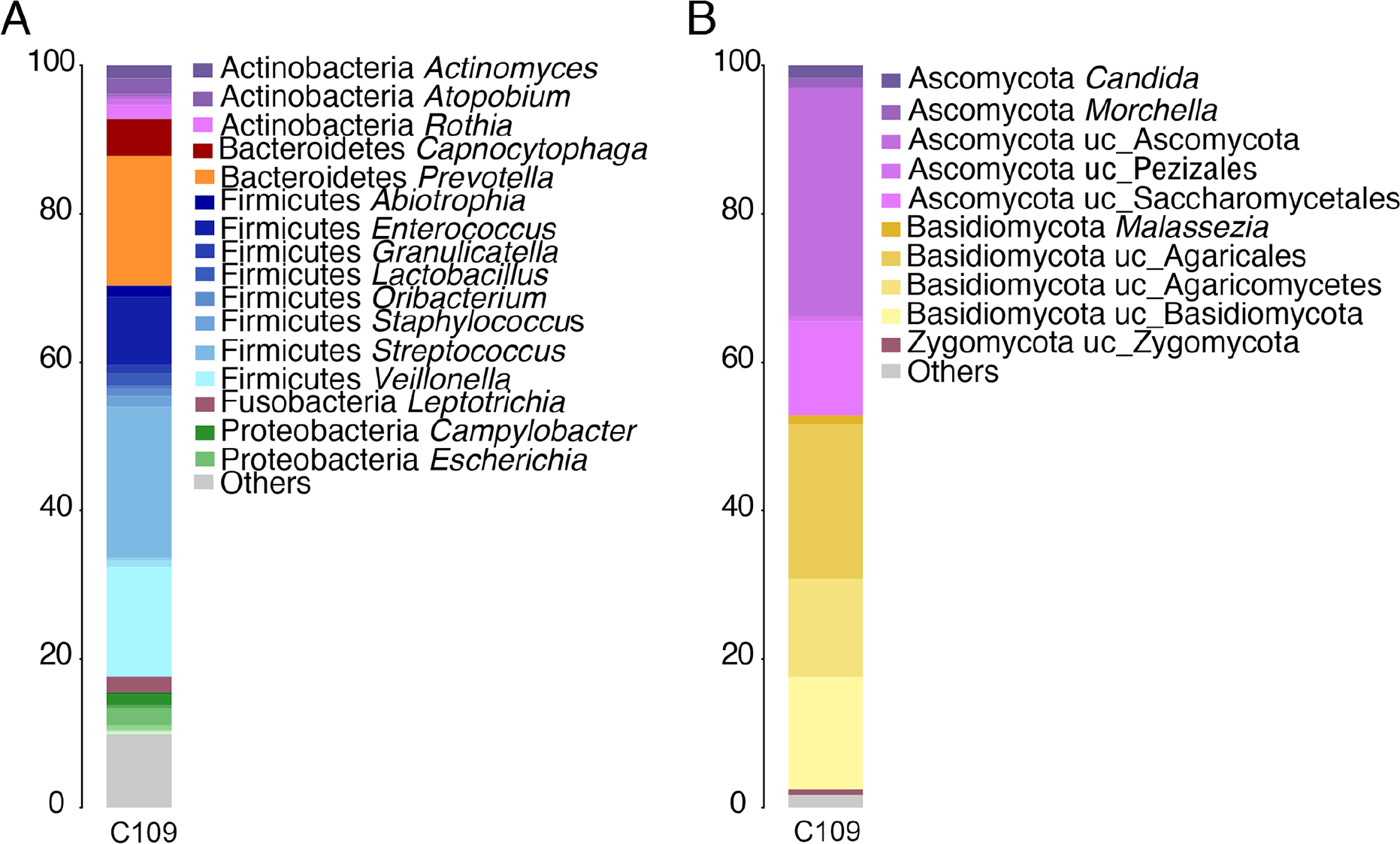
Microbiome composition of the BAL sample of patient C. **(A)** Bacterial composition. The taxonomy is based on the RDP. **(B)** Fungal composition. The taxonomy is based on the Warcup ITS training set. The phyla and genera are shown for the most abundant groups (>1%).

### Archaea are present in the lung microbiota during infection and antibiotics therapy

As it was recently reported that archaea seem to be present at all body sites including the nose and lung [34], we analysed the archaea composition in the samples of the patients of this study. We used primers 787F/1000R that amplify a 16S rRNA gene region that were specifically designed for archaea detection but that also amplify bacterial rRNA [35]. Thus, for all the samples we obtained a mixture of bacterial and archaeal reads. The proportion of bacteria and archaea was very variable between samples (**Supplementary Table 1**) as the bacterial proportion ranged from 27 to 98% reads (average of 57%), and the archaeal proportion from 2 to 72% reads (average of 43%). After filtering the bacterial reads and classification (0.5 criteria of bootstrap) in average 45% of the sequences were retained. The most abundant archaea identified in all the patients was *Methanobrevibacter* representing more than 50% of the archaeal diversity in all the samples (**Supplementary Table 2**). Patients A and C carried also other unclassified Euryarchaeota and Thermoprotei (Crenarchaeota), while in patient C only archaea belonging to the genus *Methanobrevibacter* were identified. The closest species of the most abundant OTUs was *Methanobrevibacter smithii*, which was surprising as it is described as an anaerobic methane producing archaea. It was reported to be present mainly in the human gut, however recently it was also found in the nose, lungs and on the skin [34]. Thus, it is possible that also in the lung environment there might be some anaerobic niches, or that other bacteria present in the lung help this anaerobic archaeon to grow in the lungs. Indeed, aerobic culture of methanogenic archaea without an external source of hydrogen has been reported when *Bacteroides thetaiotaomicron*, which produces hydrogen was present [36]. The presence of archea in all our samples can be related to the fact that the archaea are generally resistant to most of the antibiotics [37, 38]. Interestingly, methanogenic archaea have been associated with different diseases suggesting that this group of archaea may contribute to the disease in specific conditions and thus, perhaps also during pneumonia [34].

### *Legionella*-related amoeba were identified in the human lungs

Environmental, aquatic amoeba are the reservoir of *Legionella* and amoeba infected with *Legionella* have also been implicated as vehicle of transmission. Experimental results from a mouse model of infection support the hypothesis that inhaled protozoa may serve as cofactors in the pathogenesis of pulmonary disease induced by inhaled respiratory pathogens [39]. Thus, we analysed the composition of protozoa in the BAL samples. First standard 18S rRNA primers were used but the human contamination was too high to correctly analyse the data (data not shown). Thus, we analysed the presence of the genus *Acanthamoeba* a natural host of *Legionella* and laboratory model amoeba, and the class Heterolobosea that contains many of the known amoebal hosts of *Legionella*, with primers that are more specific for amoeba. Indeed, this allowed detecting the genus *Acanthamoeba* (**Supplementary Table 3**) and the genus *Trichomonas* in all the samples analysed (**Supplementary Table 4**). Most of the OTUs of the genus *Acanthamoeba* belonged to the species *Acanthamoeba castellani*, which is a natural host of *Legionella*. The most abundant OTU identified was close to *Trichomonas tenax* a protozoa commonly found in the human oral cavity that is also rarely associated with pulmonary infections [40]. Furthermore, a very high presence of unclassified eukaryotes was identified, suggesting that a large fraction of the eukaryotic diversity present in the lungs remains unknown (**Supplementary Tables 3** and **4**). Further studies using different genus-specific primers and samples from healthy individuals will allow to learn whether there is a community of resident protozoa in the lungs or if they are specifically associated to *Legionella* caused pneumonia.

### The lung microbiome during *Legionella* infection and antimicrobial therapy is significantly different from a healthy microbiome

To better understand the differences between the lung microbiome composition of healthy individuals and the lung microbiome influenced by infection and antibiotic treatment, we compared the healthy microbiomes reported by Segal and colleagues, 2016 [4] with our patient samples. We first reanalysed the 49 published BAL samples using our analyses pipeline. This confirmed the two groups described by the authors. The SPT cluster that is enriched in Firmicutes (Veillonellaceae, Lachnospiraceae), Bacteroidetes (Prevotellaceae, Porphyromonadaceae) and Fusobacteria (Fusobacteriaceae) and the BPT cluster that is enriched in Proteobacteria (Xanthomonadaceae, Acetobacteraceae) and Actinobacteria (Microbacteriaceae) (**Supplementary Figure 3**).

The comparison showed, that the lung microbiome composition of the healthy people and the pneumonia patients were significantly different when applying the ADONIS test (p-value=0.004975). Also, cluster analyses clustered the microbiomes of the pneumonia patient separately from the two healthy pneumotypes (**Figure 7A)**. Furthermore, the samples containing a high abundance of *Legionella* clustered together, while the two microbiomes analysed after the antibiotic treatment (one from patient B and the one from patient C) were different. However, they were also different from the healthy microbiomes.

**Figure 7.**
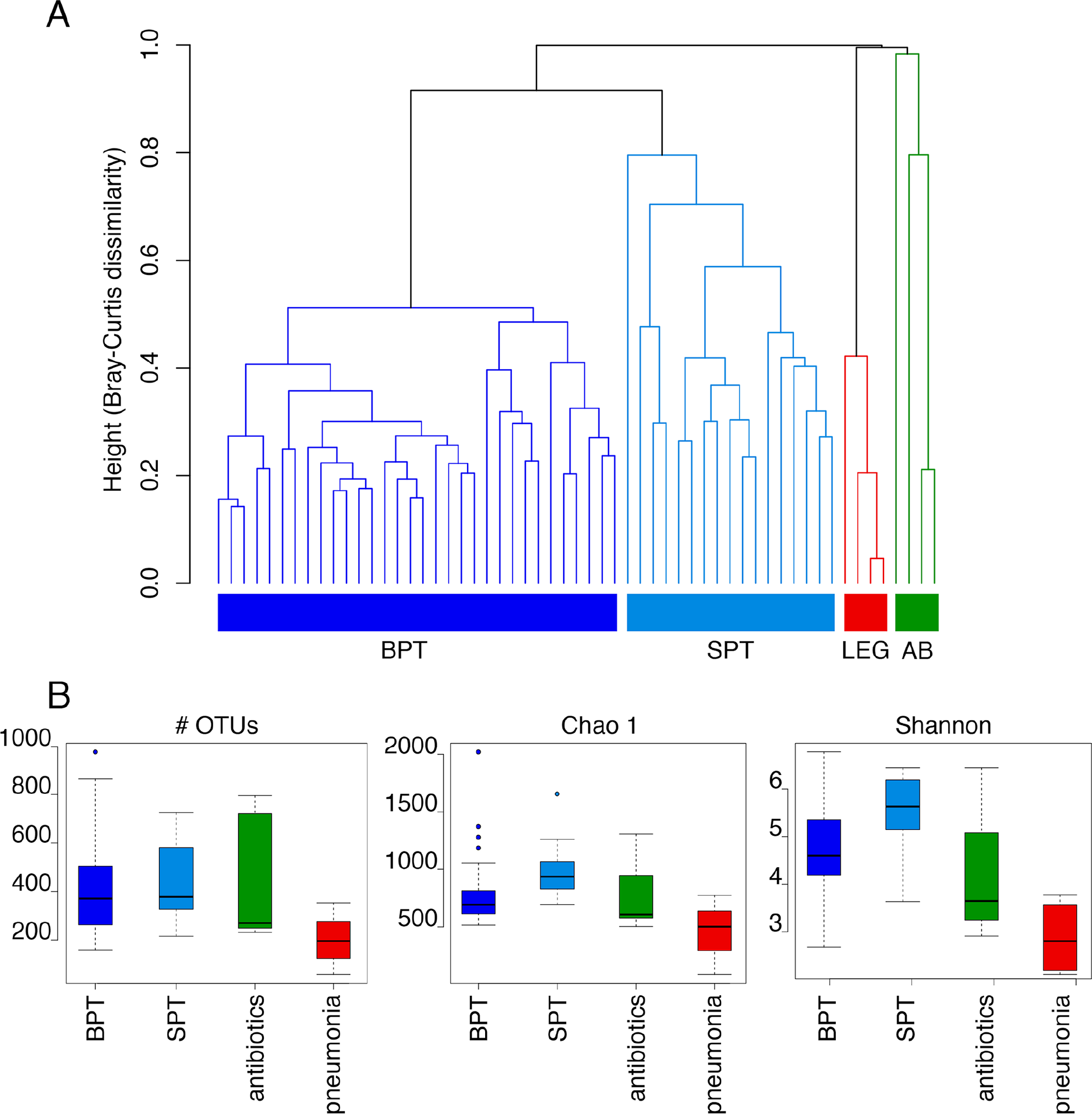
Comparison of the bacterial composition of healthy (BPT and SPT) and *Legionella*-infected and antibiotic treated BAL samples. **(A)** Hierarchical clustering of all samples based on the bacterial composition. The Bray-Curtis dissimilarity was used as distance method. (**B**) Comparison of the diversity between the two healthy pneumotypes, the pneumonia and the antibiotic treated samples. The diversity metrics Chao 1 richness estimator, number of OTUs and the Shannon Index were estimated for the three groups and statistically compared by using the Wilcoxon signed-ranked test. P-values lower than 0.05 were considered as significant.

We used the linear discriminant analysis (LDA) effect size (LEfSe) analysis and compared statistically the two pneumotypes (healthy microbiome) with the infected or the antibiotics treated samples (disrupted microbiome) (**Supplementary table 5**). This revealed that a total of 34 families of which the majority belonged to the phyla Proteobacteria, were more abundant in microbiome samples of healthy people. The most significant ones were Xanthomonadaceae, Acetobacteraceae, Sphingomonadaceae, (Proteobacteria), Veillonellaceae (Firmicutes), Flavobacteriaceae (Bacteroidetes) and Verrucomicrobiaceae (Verrucomicrobia). In contrast, bacteria belonging to these families were clearly under-represented in samples from pneumonia patients and also in those obtained after long-term antibiotic treatment. Compared to the microbiome composition of healthy lungs, a high abundance of only five families: Legionellaceae (Proteobacteria), Staphylococcaceae, Streptococcaceae (Firmicutes), Propionibacteriaceae and Corynebacteriaceae (Actinobacteria) was seen in the microbiome of the pneumonia patients. When classified at species level using BLASTN against the NCBI nucleotide database *L. pneumophila*, *Staphylococcus* sp., *Streptococcus sanguinis*, *Cutibacterium acnes* and *Corynebacterium* sp. were the most abundant OTUs from each of these families. Some species may take advantage of the host inflammatory responses that is induced by infection. For example, it has been shown that the generation of intra-alveolar catecholamines and inflammatory cytokines alters the microbial growth conditions thereby favouring specific bacterial groups during lung infections including *Streptococcus* [41]. Furthermore, a retrospective study showed that *Streptococcus sanguinis* is an important agent associated with lung abscess development (“Jerng(1997),” n.d.), a OTU that was highly abundant also in our abscess sample (40%) suggesting that this might be a general feature of lung abscess.

When the Wilcoxon signed-rank test was used to compare the diversity values between the pneumonia samples and the two pneumotypes (BPT and SPT) we found that the Chao 1 estimator was significantly lower in the pneumonia samples than in the BPT samples (p-value=0.03304) and SPT (p-value=0.001337), similar to what was seen for the number of observed OTUs compared to the BPT (p-value=0.01463) and SPT (p-value=0.01771) groups (**Figure 7B**). The Shannon index was also higher for the BPT (p-value=0.0018) and SPT (p-value=0.0006683) samples than the pneumonia microbiomes. The healthy microbiome compared to the antibiotic treated samples showed a higher abundance but not significant for the Chao 1 (BPT vs antibiotics: p-value=0.34, SPT vs antibiotics: p-value=0.14), the number of OTUs (BPT vs antibiotics: p-value=0.51, SPT vs antibiotics: p-value=0.28) and the Shannon Index (BPT vs antibiotics: p-value=0.27, SPT vs antibiotics: p-value=0.14). Thus, despite the small number of samples analysed it is clear that the bacterial species richness and evenness during pneumonia and antibiotic treatment are lower than the diversity present in healthy lungs. Low diversity has been also described in HIV-patients with acute pneumonia and for other lung conditions such as cystic fibrosis [5, 18], indicating that the overgrowth of the pathogen leads to a reduction in the microbiome diversity. However, as for the composition the antibiotics disturbed microbiome showed an intermediate diversity between pneumonia and healthy pneumotypes representing probably a “transition state”.

### Carbohydrate metabolism, glycerophospholipid and nicotinate and nicotinamide metabolism related genes are hallmarks of the pneumonia microbiome

The different composition of the lung microbiomes of the healthy and the pneumonia patients might have functional impact as different bacteria produce different metabolites and metabolise different nutrients. We thus used the software PICRUSt to predict the metagenome of the different microbiomes and compared them to identify functional differences. The LEfSe analysis was applied to statistically compare the two groups according to the functional categories represented in the KEGG pathways (**Supplementary table 6**). Indeed, 28 metabolic pathways were over-represented in pneumonia microbiomes, while 17 were more abundant in the healthy microbiomes. In the microbiome of the pneumonia patients an over-abundance of genes associated with bacterial motility such as flagella assembly or bacterial chemotaxis were identified. Furthermore, a higher abundance of genes related to carbohydrate metabolism like genes coding for the enzymes of the pentose phosphate pathway and the pentose and glucuronate interconversions was present in the microbiome during pneumonia and antibiotic therapy. This is similar to what was seen in metagenomes obtained from the human gut under antibiotic therapy where metabolic pathways related to the carbohydrate metabolism were also over-represented [42, 43]. According to gut-based studies an altered microbiome due to antimicrobial therapy releases a different repertoire of sugars, which is better exploited by opportunistic microorganisms [44–46]. The food web in the lung microbial ecosystem is far from known, but the depletion of the normal members of the microbiome seems to lead to an unbalance in the global metabolism of the community similar to what occurs in the gut, and thereby favour the growth of opportunistic microorganisms.

In the microbiome of pneumonia patients, we also identified a higher abundance of genes involved in the glycerophospholipid pathway (lipid metabolism) and the nicotinate and nicotinamide metabolism. Alterations in glycerophospholipid levels have also been found in BALs of HIV patients as well as a correlation of Staphylococacceae with a higher activity in the nicotinate and nicotinamide metabolism [47], similar to what we see in the pneumonia related microbiomes analysed here. On the other hand, the microbiome of healthy people was enriched in amino acid biosynthesis pathways. The activity of commensal bacteria in the biosynthesis and metabolism of amino acids has been described in the human gut (for a review see [48]). Certain products of the metabolism of amino acids such as short-chain fatty acids stimulate the immune system positively and have protective functions in the gut [49].

Taken together, the *Legionella* induced pneumonia altered the microbiome, leading to different metabolic activities that allow bacteria such as Staphylococacceae to become more abundant and seem to decrease bacterial activities that produce short-chain fatty acids and stimulate the immune system positively.

### Ecological interactions between bacterial and fungal communities were observed in the human lungs

Although the fungal community showed less changes than the bacterial community during antibiotic treatment, we identified some parallel changes in diversity between both groups (**Supplementary figures 1**, **2**). To get insight into the ecology of these communities and their possible interactions, we used the Wilcoxon rank-test and compared the diversity metrics distribution of both communities (**Figure 8A**). As it seen in **Figure 8A** the bacterial fraction showed a significant higher species diversity richness (Chao 1 test and number of OTUs (p-value < 0.05)), but a more equitable distribution between the different species in the fungal community was observed (p-value < 0.05). Thus, the bacterial community is more variable in its composition and richer in species than the fungal community, but the distribution is less even probably due to the dominance of the pathogen and other opportunistic bacteria during infection and antibiotic therapy. To identify whether there might be a relation between the diversity of the bacterial and the fungal communities, Pearson correlation test comparisons between the alpha-diversity metrics were performed (**Figure 8B**). A significant positive correlation of the richness (number of OTUs and Chao 1) between bacteria and fungi was identified. The species distribution (based on Shannon Index) between both communities showed a positive trend but it was not significant. This suggests a cross-domain relation between both communities and a correspondence between different bacterial and fungal species, but the relation is not strongly dependent on their abundances.

**Figure 8.**
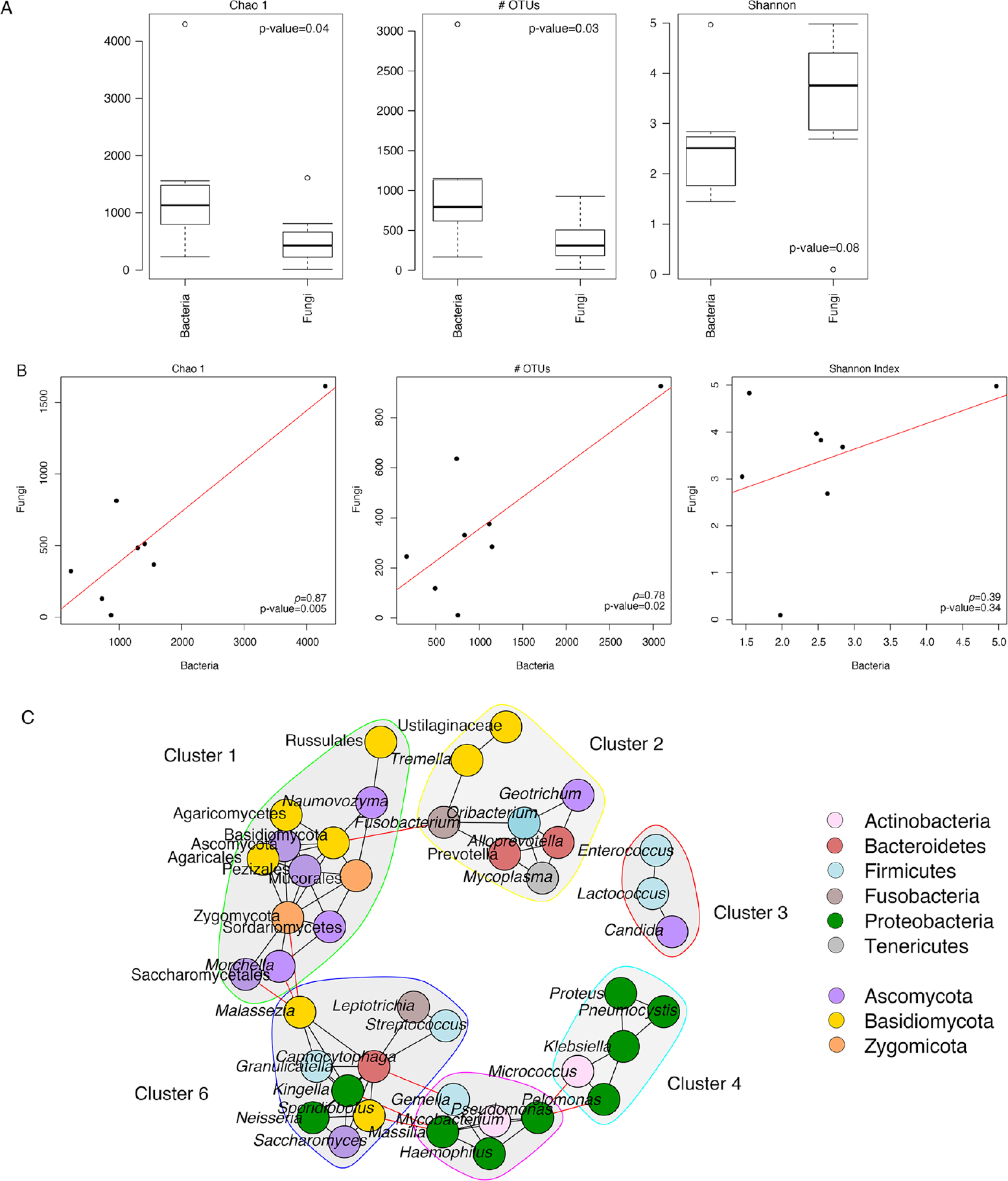
Correlation of diversity and co-occurrence network of the bacterial and fungal communities. (**A**) Comparison of the diversity between fungi and bacteria of the BAL samples. The diversity metrics Chao 1 richness estimator, number of OTUs and the Shannon Index were estimated for the two domains and statistically compared by using the wilcoxon signed-ranked test. P-values lower than 0.05 were considered as significant. (**B**) Correlation of the Chao 1 richness estimator, the number of OTUs and the Shannon Index between the bacterial and fungal communities. The correlation is based on the Pearson method and p-values < 0.05 were considered as significant. (**C**) Co-occurrence networks of bacterial and fungal communities. The network is based on the OTU-rarefied abundance tables collapsed at the genus level. Only significant positive associations are shown (p-value < 0.01). The nodes represent the different taxa involved in the network and are coloured according to their phylum.

Due to these results we established a correlation network for the microbiome and mycobiome and networks specific of each domain to identify putative associations between members of each community (**Figure 8** (bacteria-fungi), **Supplementary figure 4A** (bacteria-bacteria) and **4B** (fungi-fungi)). Interestingly, we identified bacteria-bacteria, fungi-fungi and bacteria-fungi associations, however, the relationships within each domain were the most abundant (**Figure 8**). Furthermore, certain associations identified might be very strong, as they are maintained even when we analysed all possible interactions together (bacteria-fungi). Examples are the *Mycoplasma*-*Prevotella*-*Fusobacterium* (bacteria) (**Supplementary figure 4A**) or the *Morchella*-*Malassezia*-Saccharomycetales (fungi) cluster (**Supplementary figure 4B**).

We identified six main clusters in the mixed network. Cluster 1 was composed only of fungi belonging to three different phyla, whereas cluster 2 was composed of bacteria and fungi from different phyla. Most interestingly cluster 1 and 2 were connected through a predicted interaction between *Fusobacterium* and *Basidiomycota*. For cluster 1 a second interaction with cluster 6 containing fungi and bacteria was observed through interactions between *Saccharomyces*, *Zygomycota* and *Morchella* and *Malassezia* (**Figure 8C**). Cluster 6 is further connected with the bacterial cluster 5, which shows interactions with cluster 4 containing also only bacteria. These two bacterial clusters were enriched in Proteobacteria and Actinobacteria. Cluster 3 was composed of bacteria and fungi (*Enterococcus*-*Lactococcus*-*Candida)* and is the only one for which no interactions with other clusters of the network were predicted.

To get more detailed information, which of the species are interacting, we performed the network analysis again, but based on the OTUs identified in the bacterial and fungal communities. In **Supplementary figure 5** the 30 clusters obtained are represented in a cladogram. Similarly, to the genus-based analysis, most of the clusters (25) were enriched in bacteria or fungi but 5 were composed of bacteria and fungi. We observed a trend that bacteria and fungi from the same phyla clustered. This co-occurrence of phylogenetically closer related microorganisms may suggest cooperation and/or sharing the same niche instead of competition. Associations between bacteria and fungi have been described in the literature but little is known about their mechanisms of interaction. Similar to our results where an association in a co-occurrence network of *Fusobacterium* and *Prevotella* was observed such an interaction was also described for the lung microbiome of healthy individuals [4]. Cluster 6 included the genera *Neisseria* and *Streptococcus*, both have recently been proposed as keystone species for the community structure of healthy neonatal airways [50].

Taken together, the microbial and the fungal community present in the lungs of humans are showing important interactions and seem to influence each other. *Malasezzia*, *Fusobacterium* or *Pseudomonas* seem to be key in the ecosystem since they are connecting different groups.

### Whole genome sequencing of *Legionella pneumophila* strains isolated during disease and antibiotic treatment

To follow the evolution of *L. pneumophila* during long term pneumonia and antibiotic treatment we isolated and whole genome sequenced 17 *L. pneumophila* strains at different time points over the 2-month period of disease and antibiotic treatment of patient A and 11 *L. pneumophila* strains during a 3-month period for patient B. Surprisingly, mapping of the corresponding genomes to the earliest isolate did not identify any genomic changes nor any chromosomal rearrangement (based on Pacbio assemblies) between the first and the last isolates from patient A and B. Thus *L. pneumophila* does not seem to evolve fast even when under antibiotics and immune system pressure. This is in line with our previous report where we estimated a very low evolutionary rate of 0.71 SNPs/genome/year for *L. pneumophila* strains [51].

We then searched specifically for mutations that are known to be associated with fluoroquinolones, macrolides or rifampicin resistance, as patient A had been on long term treatment with these antibiotics. However, we again did not identify any mutation. This result was intriguing, as the antibiotic therapy did not eliminate the pathogen even after 42 days of treatment. Also when the minimal extracellular concentrations inhibiting intracellular growth (MIEC) were determined for levofloxacin, erythromycin and rifampicin the *L. pneumophila* strain isolated from the pneumonia patient was not resistant to none of these antibiotics [16]. As no antibiotic resistance was found for the *L. pneumophila* strain causing the pneumonia, neither by whole genome analyses nor by susceptibility testing, the poor response to the treatment might have been due to the presence of the abscess, which has constituted a continuous source of infection. Indeed, the pneumonia for patient A was only cured after the abscess was surgically removed

## CONCLUSIONS

The analysis of the lung microbiome is still a quite new field of research but over the past decade, we have learned that the lung is not sterile, even in healthy conditions. Most analyses concentrated on lung diseases such as COPD, pulmonary cancer or cystic fibrosis. Thus, the impact of the lung microbiome on disease development and outcome of bacterial pneumonia in non-cystic fibrosis patients is nearly not known. Very recently the analyses of the lung microbial community of adenovirus pneumonia, *M. pneumoniae* pneumonia and tracheomalacia showed that each differed significantly with the dominance of the pathogen within the microbiome [52]. Our comprehensive, longitudinal analyses of the lung microbiome (bacteria, fungi, archaea and eukaryotes) of patients suffering for several month from pneumonia due to *L. pneumophila*, and its comparison with healthy lung microbiomes showed that infections engenders a highly disturbed lung microbiome. It is characterized by a low species diversity, low abundance of many different taxa (specially from the phyla Proteobacteria and Bacteroidetes) and the over-abundance of pathogenic and/or opportunistic bacteria associated with lung infections such as *Staphylococcus* sp., *Streptococcus sanguinis*, *Cutibacterium acnes* and *Corynebacterium sp*.. The strong dominance of the pathogen overgrows and outcompetes the normal, resident microbiota. The low abundance of many of these “healthy families” may be related to their susceptibility to antibiotics or to the fact that these protective and/or beneficial microorganisms were absent or lowly abundant in the patient’s microbiome before infection. Similarly, to what has been described in other body habitats, the antibiotic treated microbiome does not represent a healthy lung microbiome and shows a higher abundance of resistant bacteria such as *Staphylococcus* or *Enterococcus*.

Very little is known about the role of fungi in the health of the lungs and the response to infection. Here we show that the mycobiome follows a more stable dynamics during infection and antibiotic treatment with Ascomycota and Basidiomycota as the most abundant phyla. The “gap” in the fungi databases does not allow an accurate classification of fungi at low taxonomic levels. Thus, a great portion of the diversity at genus and family levels remains unknown so far. The identification of correlations in the diversity and composition of bacterial and fungal communities in the lungs of the here analysed individuals may suggest an ecological relationship between both communities that could be key for the restoration of the microbiome after the cessation of the disease and the antimicrobial therapy.

Interestingly, archaea might also play an important role in the lung microbiota community and in disease severity or outcome, as suggested by the presence of several different archaea and in particular of *Methanobrevibacter smithii* in all our lung microbiomes.Corrections version 6 Furthermore, we showed here that amoeba and other protozoa are present in the lung microbiome. This might partly be due to the fact that *Legionella-*infected amoeba had been inhaled we identified several different protozoa, thus it seems that the lung microbiota also contains a community of protozoa. Whether protozoa are part of the healthy microbiome or are only present in *Legionella* associated pneumonia patients, needs to be studied further.

Recently, a 3D model of human lungs was developed and the microbiome and metabolome obtained from a cystic fibrosis lung was mapped [53]. This study showed how the microorganisms occupy different regions of the lung, and how drugs penetrate the lungs at many different levels. Thus, the response of the microbiome to infection and the antibiotics depends on many factors, including the microbial composition before the treatment, the type of antibiotic and its pharmacology properties (time of retention, doses, diffusion, etc), the immunological status of the individual and the repertoire of resistance genes in the microbiome. Here we show that also interactions between fungi, archaea and protozoa need to be considered in the outcome of the infection and antibiotic therapy.

## MATERIAL AND METHODS

### Sample collection, *Legionella* detection and DNA extraction

Bronchoalveolar lavages (BAL), sputum, and a biopsy from the lung abscess were collected and stored at −80º until further processing. DNA was extracted from 1 ml of BAL sample using the PowerSoil® DNA Isolation Kit (Mobio) following the manufacturer’s instructions. The presence of *L. pneumophila* in the samples was detected by diagnostic PCR using primers and probes of the R-DiaLegTM kit (Diagenode, Belgium). The PCRs were prepared in a final volume of 20 μL by adding the Master Mix 5X (TaqMan Probe LC2.0, Roche Diagnostics, France), 4 μL of primers and probe Lspp&Lp (R-DiaLegTM) or the internal control (DICD-YDL100) 2 μL, H2O (PCR grade) 2 μL and 10 μL of the extracted sample. The amplification was performed on a LC2 system using the following program: 95◦ C for 10 min followed by 45 cycles of 95◦C for 10 s, 60◦C for 40 s and 72◦C for 1 s and a final step of 30 s at 40◦C.

### Whole genome sequencing

To analyse the intra-patient evolution of *L. pneumophila*, a total of 17 isolates from patient A and 11 from patient B during different stages of the antibiotic treatment were collected and sequenced. The isolates from patient A were taken at days 4, 5, 10, 13, 14 (2 isolates), 16, 22, 26, 29, 30, 31, 33, 34, 36, 39 and 42 during hospitalization and the isolates of patient B were taken at days 0 (6 isolates) and 69 (5 isolates) and sequenced on a Illumina Nextseq500 (150 bp paired-end) machine. Reads were trimmed for low quality and adapter removing using trimmomatic 0.36 [54]. Trimmed reads were assembled using SPAdes [55]. Polymorphisms between isolates from the same Patient were searched by mapping the trimmed reads on the assembly of the 1^st^ isolate using BWA v0.7.12 [56]. Pacbio sequencing was performed by GATC Biotech. Genomes were assembled using the SMRT Analysis software suite.

Mutations in the genes *rplD* and *rpIV* encoding ribosomal proteins, as well as in the domain V of 23S rRNA where mutations associated with macrolide resistances may occur, were analysed. Possible mutations associated to fluoroquinolone resistance in the genes *gyrA*, *gyrB* (topoisomerase II) and *parC* (topoisomerase IV) and mutations conferring rifampicin resistance in the *rpoB* gene encoding for the β subunit of the RNA polymerase were searched. Trimmed reads were mapped on the different gene targets using BWA v0.7.12 and variant calling was done using the Naïve variant caller v0.0.4 [56].

### Microbiome sequencing

For characterizing the bacterial fraction, the V3-V4 region of the 16S rRNA gene was amplified by PCR using the 16SrRNA Illumina sequencing standard primers with adapters. Forward primer (5’-CCTACGGGNGGCWGCAG-3’) with adaptor (5’-TCGTCGGCAGCGTCAGATGTGTA TAAGAGACAG-3)’ and reverse primer (5′-GACTACHVGGGTATCTAATCC-3’) with adaptor (5’-GTCTCGTGGGCTCGGAGATG TGTATAAGAGACAG-3’). For each sample a 20 μl PCR mix was prepared, containing 5 μl of Buffer Taq (10X), 1 μl of 25 mM MgCl2, 0.5 μl of dNTPs (10 mM), 1.25 μl of each primer (10 mM), 0.25 μl of Phusion High-Fidelity DNA Polymerase (5u/μl), 0.5 μl of DMSO, 8.25 μl of nuclease-free water and 1 μl of DNA template. PCR conditions were: 95◦ C for 5 min followed by 25 cycles of 95◦C for 30 s, 55◦C for 1 min and 72◦C for 1 min and a final extension step of 7 minutes at 72◦C.

For characterizing the mycobiome we amplified the ITS region by using the ITS1/ITs2 primers. Forward primer (5’-CTTGGTCATTTAGAGGAAGTAA-3’) with adaptor (5’-TCGTCGGCAGCGTCAGATGTGTATAAGAGACAG-3’) and reverse primer (5’-GCTGCGTTCTTCATCGATGC-3’) with adaptor (5’-GTCTCGTGGGCTCGGAG ATGTGTATAAGAGACAG-3’). We used the same reaction mix and PCR conditions as described above, but 30 cycles. All the amplicons were checked by electrophoresis in agarose gel (1.4%).

To characterize the archaeal microbiome a commonly used region of the 16S rRNA to detect archaea was amplified by using the primers: 787F (5’-ATTAGATACCCSBGTAGTCC-3’) with and 1000R (5’-GGCCATGCACYWCYTCTC-3’). The same sequencing adaptors and reaction mix described above were used for the PCR with the conditions: 95◦ C for 5 min followed by 30 cycles of 95◦C for 30 s, 62◦C for 1 min and 72◦C for 1 min and a final extension step of 7 minutes at 72◦C.

Two pair of primers to analyse the amoeba composition were: Primers JDP1 (5’-GGCCCAGATCGTTTACCGTGAA-3’) and JDP2 (5’-TCTCACAAGCTGCTAGGGAG TCA-3’) [57], and Vahl730F_C (5’-TAATACTGCTGTAGTTAAAACGCCC-3’) [58] and R-1200 (CCCGTGTTGAGTCAAATTAAGC) [59]. The same sequencing adaptors and reaction mix described above were used for the PCR with the conditions: 95◦ C for 5 min followed by 30 cycles of 95◦C for 30 s, 61◦C for 1 min and 72◦C for 1 min and a final extension step of 7 minutes at 72◦C for the primers 2F/12R. The same conditions were used for the other primers except for the annealing temperatures, being 63º and 57º for the JD1/JD2 and Vahl730F_C/ R-1200, respectively. For all PCR reactions negative controls were included. The Illumina libraries were then prepared following the manufacturer’s instructions. High-throughput sequencing of the amplicons was performed with a Miseq Illumina sequencer (2×300 bp) by the Biomics Pole (Institute Pasteur).

### Bacterial density determination

Quantitative real-time PCR (qPCR) was performed to quantify total bacteria and fungi in the samples based on the 16SrRNA gene. We used the forward primer 520F (5’-AYTGGGYDTAAAGNG-3’) and the reverse primer 802R (5’-TACNVGGGTATCTAATCC-3’). Standard curves were estimated by using serial 10-fold dilutions of purified and quantified amplicons. The PCR was prepared in a final volume of 20 μL by adding 10 μL of the SYBR Green QPCR Master Mix 5X (Applied Biosystems), 0.8 μL of primers (10 mM), 3.4 μL of H2O and 5 μL of the DNA sample. The amplification was performed on a Bio-Rad CFX qPCR Instrument using the following program: 95◦ C for 3 min followed by 40 cycles of 95◦C for 5 s, 55◦C for 30 s and 72◦C for 1 s. All reactions including negative controls were run in triplicates.

### Analysis of the amplicon sequences

Artefactual sequences, short reads (<50 bp), as well as sequences of low quality (Q < 33) were discarded by using scripts (fastx_artifacts_filter and fastq_quality_trimmer) from the FASTX-Toolkit (http://hannonlab.cshl.edu/fastx_toolkit/index.html). The trimming of the sequences according to quality parameters was based on the PRINSEQ (http://prinseq.sourceforge.net). Paired-end reads were joined by using the fastq-join script (http://code.google.com/p/ea-utils). Quantitative Insights Into Microbial Ecology (QIIME) pipeline was used to discard chimerical sequences and to calculate the Operational taxonomic units (OTUs) at 97% of similarity [60]. We selected the open-reference OTU picking method with the QIIME default taxonomy-training database. The taxonomic classification of the 16SrRNA and 18S rRNA reads was based on the Ribosomal Database Project (RDP) [61] and the ITS taxonomy on the Warcup ITS training set [62]. Annotations with confidence values greater than 0.5 for bacteria, archaea and eukaryotes and 0.25 for fungi were included, setting the assignment of the reads to the last-well identified taxonomic level.

### Prediction of functional features

The software Phylogenetic investigation of communities by Reconstruction of Unobserved States (PICRUSt) to computationally predict the genomic potential (virtual metagenome) of the samples from the 16S rRNA data was applied [63]. The program uses the information available for the marker gene to predict gene family abundances. We used the GreenGenes database (version 13.5) implemented in the Galaxy Server (http://huttenhower.sph.harvard.edu/galaxy/) for the prediction and the KEGG Orthologs annotation for the output.

### Ecological and statistical analysis

We estimated the microbial richness and diversity of the samples by calculating the total number of OTUs, the Chao 1 estimator [64], and the Shannon diversity index (Shannon, n.d.). All these analyses were based on the core diversity analysis script implemented in the QIIME pipeline using the same depth for all the samples. The 16SrRNA abundance table was rarefied at 60,000 reads per sample and the ITS at 6,000 according to the sequencing effort. A hierarchical clustering analysis based on Bray–Curtis dissimilarity was also applied to compare the microbial composition between the different samples.

To statistically compare the diversity estimators between groups of samples, we used the Wilcoxon signed-rank test. Correlation analyses of the diversity and composition of the microbiome and mycobiome were applied based on the Pearson method. We used a multivariate ANOVA based on dissimilarities (Adonis) to test the influence of external variables in explaining differences in composition between different groups of samples. We used the Vegan package of the R program for this analysis [65]. All analyses were based on the R software considering p-values below 0.05 as significant [66]. The linear discriminant analysis (LDA) effect size (LEfSe) analysis was applied to identify those taxa and genes characterizing the different groups of samples [67]. We considered taxa as discriminative of groups when the LDA score was higher than 2.

### Co-occurrence network analysis

We calculated co-occurrence networks for the 16SrRNA and the ITS OTU data. The networks were based on the OTU-rarefied abundance tables collapsed at genus level. We used SparCC to estimate the correlation coefficients between the genera, using 500 bootstraps from the genus-table to estimate the "two-sided" p-values [68]. The correlation matrix was filtered to keep significant correlations based on a p-value lower than 0.01 and converted in a network by using the igraph package implemented in R software [69]. The igraph package was used to display the network using a force-directed layout, representing the edges the correlation coefficients between taxa. We detected groups of taxa more connected between then and with fewer connections with other groups by using the Newman-Girvan algorithm also implemented in the igraph package.

## Ethics approval and consent to participate

The clinical sample collection of the Lyon University Hospital was declared to the French Ministry of Education and Research (number DC-2008-176) and written informed consent from patients have been obtained for this study

## Availability of data and material

All sequences have been entered in the European Bioinformatics Institute database, under project accession number: PRJEB33790.

## Authors’ contributions

AEPC and CG contributed to DNA extraction, library preparation and sequencing, AEPC and CR to data analyses and interpretation, SJ contributed to sample collection and study design. The manuscript was written by AEPC and CB with input from co-authors. The project was conceived, planned and supervised by CB.

## Acknowledgements

We thank the Institut Pasteur Biomics pole (Christiane Bouchier and Laurence Ma) for sequencing of the samples reported in this study.

