## Supplemental material for "*Legionella pneumophila* infection and antibiotic treatment engenders a highly disturbed pulmonary microbiome with decreased microbial diversity"

**Supplementary table 1**: Percentage of reads classified as bacteria or archaea for each sample

**Supplementary table 2**: Relative abundance of archaea present in the BAL samples

**Supplementary table 3**: Relative abundance of protozoa in the BAL samples

**Supplementary table 4**. Relative abundance of eukaryotic sequences in the BAL samples

**Supplementary table 5**. Bacterial families identified with statistically significantly different abundance between healthy and patient samples

**Supplementary table 6**. KEGG pathways identified having different abundances between samples of healthy people and the patient samples

**Supplementary figure 1**. Bacterial alpha-diversity of patient A and B

**Supplementary figure 2**. Fungal alpha-diversity of patient A and B

**Supplementary figure 3**. Lung microbiome composition of healthy (SPT and BPT) and pneumonia samples

**Supplementary figure 4**. Correlation of taxa in lung microbiome during pneumonia

**Supplementary figure 5**. Co-occurrence network of bacterial and fungal communities

**Supplementary table 1**. Percentage of reads classified as bacteria or archaea for each sample.

| Sample | Bacteria (%) | Archaea (%) | Archaeal reads classified at bootstrap 0.5 |
|---|---|---|---|
| PatA-day5 | 27.3 | 72.7 | 6.7 |
| PatA-day14 | 44.1 | 55.9 | 2.7 |
| PatA-day24 | 36.3 | 63.7 | 12.6 |
| PatA-day33 | 34.8 | 65.2 | 10.7 |
| PatA-day42 | 98.3 | 1.7 | 97 |
| PatB-day0 | 89.9 | 10.1 | 89 |
| PatB-day82 | 92.3 | 7.7 | 91 |
| PatC-day109 | 35.9 | 64.1 | 52.4 |
| **average** | 57.3 | 42.7 | 45.3 |

The proportion of archaeal reads was calculated based on RDP taxonomic assignation. Results for considering a minimum criterium of bootstraps of 0.5 arealso shown. PatA5, PatA14, PatA33, PatA42, BAL sample of patient A at day 5, 14, 33, and 42;PatB0, PatB82, BAL sample of Patient B at day 0 and 82; PatC109, BAL sample of patient C at day 109

**Supplementary table 2**. Relative abundance of archaea present in the BAL samples

| Taxonomy | PatA5 | PatA14 | PatA24 | PatA33 | PatA42 | PatB0 | PatB82 | PatC109 |
|---|---|---|---|---|---|---|---|---|
| Crenarchaeota;uc_Thermoproteaceae | 8.8 | 4.8 | 5.8 | 1.6 | 0 | 1.3 | 8.8 | 0 |
| Crenarchaeota;uc_Thermoproteales | 0.4 | 0.4 | 0 | 0 | 0 | 0 | 0.4 | 0 |
| Crenarchaeota;uc_Thermoprotei | 13.9 | 18.1 | 14.4 | 2.5 | 5.6 | 1.7 | 13.9 | 0 |
| Euryarchaeota;Methanobrevibacter | 73.8 | 51.5 | 56.9 | 70.7 | 55.6 | 95.6 | 73.8 | 100 |
| Euryarchaeota;Methanoregula | 0 | 4 | 0 | 0 | 0 | 0 | 0 | 0 |
| Euryarchaeota;uc_Euryarchaeota | 3.0 | 21.1 | 22.9 | 25.2 | 38.9 | 1.3 | 3.0 | 0 |

Classification of archaeal reads was based on RDP taxonomy. PatA5, PatA14, PatA33, PatA42, BAL sample of patient A at day 5, 14, 33, and 42;PatB0, PatB82, BAL sample of Patient B at day 0 and 82; PatC109, BAL sample of patient C at day 109

**Supplementary table 3**. Relative abundance of protozoa in the BAL samples

| Taxonomy | PatA5 | PatA14 | PatA24 | PatA33 | PatA42 | PatB0 | PatB82 | PatC109 |
|---|---|---|---|---|---|---|---|---|
| Eukaryota;uc_Acanthamoeba | 52.0 | 28.5 | 35.1 | 44.3 | 47.6 | 36.4 | 44.6 | 39.8 |
| Eukaryota;uc_Bilateria | 0.9 | 2.6 | 2.9 | 1.6 | 1.6 | 1.1 | 0.4 | 1.7 |
| Eukaryota;uc_Opisthokonta | 0.0 | 1.7 | 0.0 | 0.0 | 0.0 | 0.0 | 0.0 | 0.0 |
| Eukaryota;uc_Eukaryota | 47.1 | 67.1 | 62.0 | 54.1 | 50.7 | 62.5 | 55.1 | 58.5 |

Amoeba primers used were JDP1/JDP2. The classification was based on SILVA database (SILVA_132_QIIME_release). PatA5, PatA14, PatA33, PatA42, BAL sample of patient A at day 5, 14, 33, and 42;PatB0, PatB82, BAL sample of Patient B at day 0 and 82; PatC109, BAL sample of patient C at day 109

**Supplementary table 4**. Relative abundance of eukaryotic sequences in the BAL samples

| Taxonomy | PatA5 | PatA14 | PatA24 | PatA33 | PatA42 | PatB0 | PatB82 | PatC109 |
|---|---|---|---|---|---|---|---|---|
| Eukaryota;Trichomonas | 1.4E-02 | 0.3 | 89.4 | 70 | 0.7 | 66.7 | 88.1 | 0.9 |
| Eukaryota;uc_Bilateria | 0.6 | 0.7 | 0.1 | 0.2 | 6.8 | 0.5 | 0.9 | 4.3 |
| Eukaryota;uc_Eumetazoa | 0.1 | 0 | 0 | 3,8E-02 | 0.1 | 0 | 0.1 | 0.1 |
| Eukaryota;uc_Opisthokonta | 0.4 | 13.8 | 1.0 | 5.2 | 8.3 | 2.2 | 0.5 | 6.4 |
| Eukaryota;uc_Eukaryota | 64.9 | 85.0 | 9.4 | 24.2 | 84.2 | 30.5 | 10.4 | 88.3 |
| Unassigned | 34.0 | 0.1 | 0 | 0.3 | 0 | 0.1 | 0 | 0.1 |

Primers Vahl730F_C/R-1200 were used. The classification is based on SILVA database (SILVA_132_QIIME_release). PatA5, PatA14, PatA33, PatA42, BAL sample of patient A at day 5, 14, 33, and 42;PatB0, PatB82, BAL sample of Patient B at day 0 and 82; PatC109, BAL sample of patient C at day 109

**Supplementary table 5**. Bacterial families identified with statistically significantly different abundance between healthy and patient samples.

| Family | Sample type | LDA score | P-value |
|---|---|---|---|
| Xanthomonadaceae | healthy | 5.7 | 6.68E-06 |
| Veillonellaceae | healthy | 4.86 | 1.40E-03 |
| Flavobacteriaceae | healthy | 4.58 | 1.15E-02 |
| Acetobacteraceae | healthy | 4.52 | 1.17E-05 |
| Sphingomonadaceae | healthy | 4.28 | 2.22E-05 |
| Oxalobacteraceae | healthy | 4.24 | 1.57E-05 |
| Porphyromonadaceae | healthy | 4.14 | 1.58E-03 |
| Verrucomicrobiaceae | healthy | 4.12 | 2.21E-06 |
| Pseudomonadaceae | healthy | 4.08 | 6.46E-03 |
| Moraxellaceae | healthy | 4.07 | 2.42E-02 |
| Cellulomonadaceae | healthy | 4.03 | 1.95E-05 |
| Comamonadaceae | healthy | 3.86 | 1.14E-03 |
| Planctomycetaceae | healthy | 3.84 | 9.89E-06 |
| Sphingobacteriaceae | healthy | 3.83 | 5.46E-05 |
| Microbacteriaceae | healthy | 3.82 | 6.99E-05 |
| Chitinophagaceae | healthy | 3.76 | 8.87E-05 |
| Ruminococcaceae | healthy | 3.43 | 5.22E-04 |
| Cytophagaceae | healthy | 3.40 | 1.14E-04 |
| Burkholderiaceae | healthy | 3.38 | 1.40E-05 |
| Rhodobacteraceae | healthy | 3.34 | 1.72E-02 |
| Rhodocyclaceae | healthy | 3.30 | 3.26E-05 |
| Nocardiaceae | healthy | 3.23 | 2.53E-02 |
| Erythrobacteraceae | healthy | 3.17 | 7.75E-03 |
| Parachlamydiaceae | healthy | 3.17 | 1.64E-02 |
| Beijerinckiaceae | healthy | 3.16 | 1.23E-03 |
| Clostridiaceae | healthy | 3.10 | 2.75E-03 |
| Methylophilaceae | healthy | 3.09 | 2.12E-03 |
| Bdellovibrionaceae | healthy | 3.04 | 7.27E-03 |
| Bacteriovoracaceae | healthy | 3.03 | 1.13E-02 |

**Supplementary table 6**. KEGG pathways identified having different abundances between samples of healthy people and the patient samples.

| Category | type | LDA | p-value | Role |
|---|---|---|---|---|
| Pentose phosphate pathway | pneumonia | 2.86 | 1.E-04 | Carbohydrate Metabolism |
| Pentose and glucuronate interconversions | pneumonia | 2.65 | 8.E-04 | Carbohydrate Metabolism |
| Pyruvate metabolism | pneumonia | 2.55 | 4.E-02 | Carbohydrate Metabolism |
| Bacterial motility proteins | pneumonia | 3.04 | 7.E-04 | Cell Motility |
| Flagellar assembly | pneumonia | 2.56 | 1.E-02 | Cell Motility |
| Bacterial chemotaxis | pneumonia | 2.2 | 4.E-02 | Cell Motility |
| Other transporters | pneumonia | 2.68 | 9.E-04 | Cellular Processes and Signalling |
| Other ion-coupled transporters | pneumonia | 2.27 | 4.E-02 | Cellular Processes and Signalling |
| Peptidases | pneumonia | 2.87 | 1.E-02 | Enzyme Families |
| Protein processing in endoplasmic reticulum | pneumonia | 2.47 | 1.E-03 | Folding |
| Translation proteins | pneumonia | 2.87 | 8.E-03 | Genetic Information Processing |
| Transcription factors | pneumonia | 2.74 | 5.E-03 | Genetic Information Processing |
| Biosynthesis of unsaturated fatty acids | pneumonia | 2.63 | 1.E-02 | Lipid Metabolism |
| Fatty acid metabolism | pneumonia | 2.52 | 5.E-02 | Lipid Metabolism |
| Glycerophospholipid metabolism | pneumonia | 2.33 | 4.E-02 | Lipid Metabolism |
| Nicotinate and nicotinamide metabolism | pneumonia | 2.97 | 7.E-05 | Metabolism of Cofactors and Vitamins |
| Thiamine metabolism | pneumonia | 2.73 | 4.E-05 | Metabolism of Cofactors and Vitamins |
| Vitamin B6 metabolism | pneumonia | 2.62 | 9.E-04 | Metabolism of Cofactors and Vitamins |
| Biotin metabolism | pneumonia | 2.63 | 3.E-04 | Metabolism of Cofactors and Vitamins |
| Riboflavin metabolism | pneumonia | 2.24 | 7.E-03 | Metabolism of Cofactors and Vitamins |
| Taurine and hypotaurine metabolism | pneumonia | 2.25 | 4.E-02 | Metabolism of Other Amino Acids |
| Tetracycline biosynthesis | pneumonia | 2.473 | 5.E-02 | Metabolism of Terpenoids and Polyketides |
| Purine metabolism | pneumonia | 2.78 | 3.E-02 | Nucleotide Metabolism |
| Function unknown | pneumonia | 2.96 | 1.E-02 | Poorly Characterized |
| Chromosome | pneumonia | 3.17 | 9.E-03 | Replication and Repair |
| Drug metabolism other enzymes | pneumonia | 2.83 | 2.E-03 | Xenobiotics Biodegradation and Metabolism |
| Benzoate degradation | pneumonia | 2.44 | 3.E-04 | Xenobiotics Biodegradation and Metabolism |
| Aminobenzoate degradation | pneumonia | 2.41 | 6.E-03 | Xenobiotics Biodegradation and Metabolism |
| Glycine, serine and threonine metabolism | health | 3.07 | 2.E-03 | Amino Acid Metabolism |
| Lysine biosynthesis | health | 2.96 | 7.E-04 | Amino Acid Metabolism |
| Valine, leucine and isoleucine biosynthesis | health | 2.89 | 8.E-03 | Amino Acid Metabolism |
| Cysteine and methionine metabolism | health | 2.70 | 2.E-02 | Amino Acid Metabolism |
| Novobiocin biosynthesis | health | 2.30 | 5.E-02 | Biosynthesis of Other Secondary Metabolites |
| C5-Branched dibasic acid metabolism | health | 2.73 | 3.E-03 | Carbohydrate Metabolism |
| Inositol phosphate metabolism | health | 2.43 | 4.E-03 | Carbohydrate Metabolism |
| Cell motility and secretion | health | 2.42 | 3.E-04 | Cellular Processes and Signalling |
| Oxidative phosphorylation | health | 3.23 | 2.E-02 | Energy Metabolism |
| Carbon fixation in photosynthetic organisms | health | 2.57 | 6.E-03 | Energy Metabolism |
| Protein export | health | 2.93 | 6.E-04 | Folding |
| RNA transport | health | 2.22 | 5.E-02 | Translation |
| One carbon pool by folate | health | 2.62 | 5.E-03 | Metabolism of Cofactors and Vitamins |
| Lipoic acid metabolism | health | 2.31 | 2.E-03 | Metabolism of Cofactors and Vitamins |
| Prenyltransferases | health | 2.57 | 5.E-03 | Metabolism of Terpenoids and Polyketides |
| Cell cycle, Caulobacter | health | 2.66 | 4.E-03 | NA |
| Ubiquinone and terpenoid, quinone biosynthesis | health | 2.66 | 4.E-03 | NA |

Linear discriminant analysis (LDA) scores higher than 2 and p-values lower than 0.05 were considered as significant.
The main functions encoded by the different pathways are shown in the last column

| | | | |
|---|---|---|---|
| Opitutaceae | healthy | 3.94 | 2.91E-03 |
| Rhodospirillaceae | healthy | 2.61 | 1.62E-02 |
| Oceanospirillaceae | healthy | 2.58 | 5.76E-04 |
| Gallionellaceae | healthy | 2.50 | 3.56E-03 |
| Micromonosporaceae | healthy | 2.48 | 2.45E-02 |
| Legionellaceae | pneumonia | 5.66 | 8.96E-14 |
| Staphylococcaceae | pneumonia | 4.63 | 4.67E-06 |
| Streptococcaceae | pneumonia | 4.59 | 8.71E-06 |
| Propionibacteriaceae | pneumonia | 3.49 | 2.39E-06 |
| Corynebacteriaceae | pneumonia | 3.44 | 2.02E-05 |

Linear discriminant analysis (LDA) scores higher than 2 and p-values lower than 0.05 were considered as significant

### Figure Legends

**Supplementary figure 1**. **Bacterial alpha-diversity of patient A and B**. Bacterial diversity was based on the 16SrRNA OTU-table. The diversity metrics Chao 1 richness estimator, number of OTUs and the Shannon Index were calculated. (**A**) Patient A, (**B**) Patient B.

**Supplementary figure 2**. **Fungal alpha-diversity of patient A and B**. Fungal diversity was based on the ITS OTU-table. The diversity metrics Chao 1 richness estimator, number of OTUs and the Shannon Index were calculated. (**A**) Patient A,Figure preparation (**B**) Patient B.

**Supplementary figure 3**. **Lung microbiome composition of healthy (SPT and BPT) and pneumonia samples**. The taxonomy is based on the RDP. The phyla and genera are shown for the most abundant groups (>2.5%).

**Supplementary figure 4**. **Correlation of taxa in lung microbiome during pneumonia**. (**A**) Correlation of bacterial genera (**B**) Correlation of fungal genera. The networks are based on the OTU-rarefied abundance tables collapsed at genus level. Only significant positive associations are shown (p-value < 0.01). The nodes represent the different taxa involved in the networks and are coloured by phylum

**Supplementary figure 5**. **Co-occurrence network of bacterial and fungal communities**. The network is based on the OTU-rarefied abundance tables. Only significant positive associations are shown (p-value < 0.01). The network is represented as a cluster. Each branch represents an OTU involved in the network and are coloured by phylum. The figure was divided in three blocks for a better visualization of the details.

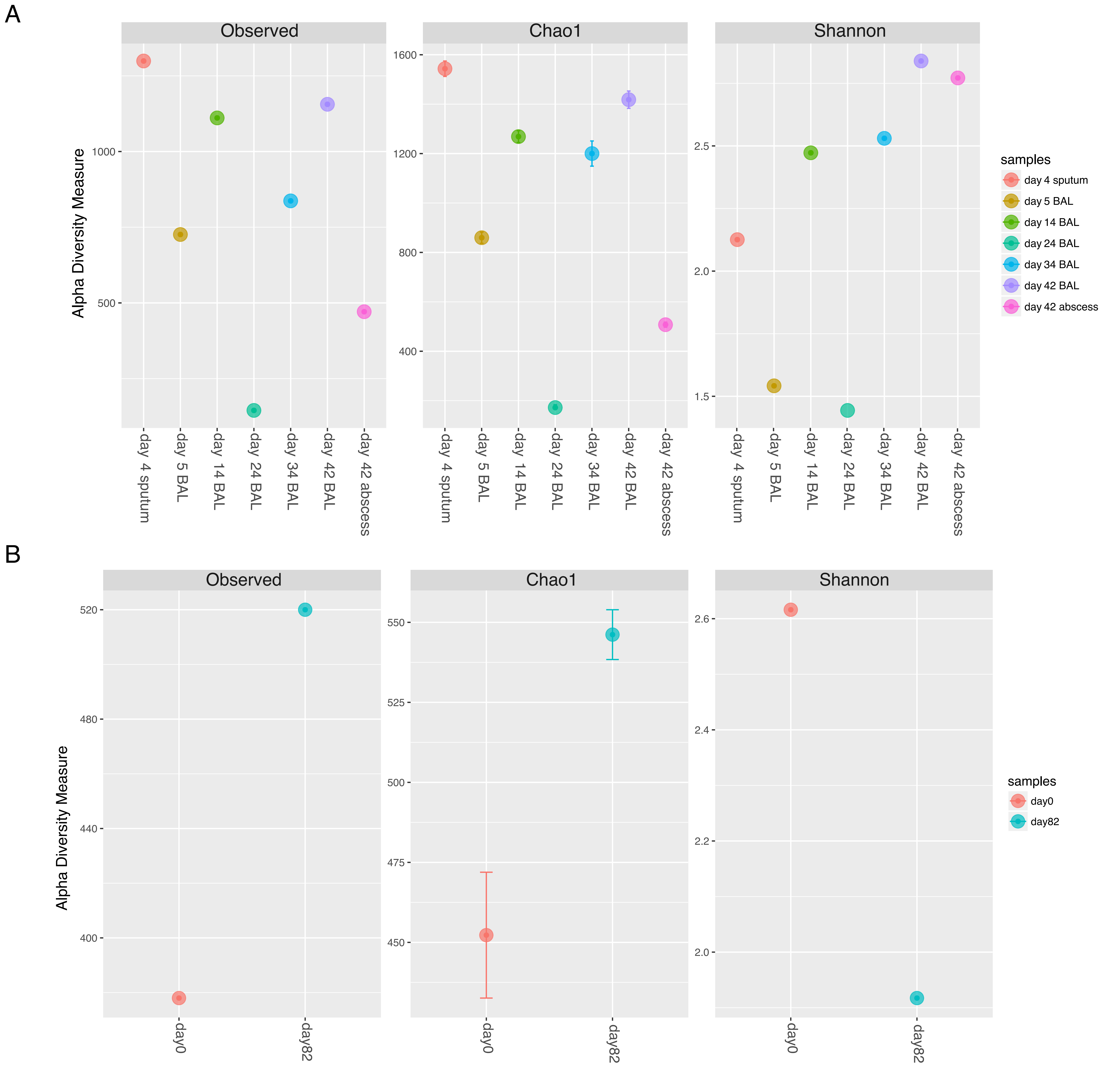
A
Observed
Chao1
Shannon
Alpha Diversity Measure
1000
500
1600
1200
800
400
2.5
2.0
1.5
day 4 sputum
day 5 BAL
day 14 BAL
day 24 BAL
day 34 BAL
day 42 BAL
day 42 abscess
samples
B
520
480
440
400
550
525
500
475
450
2.6
2.4
2.2
2.0
day0
day82

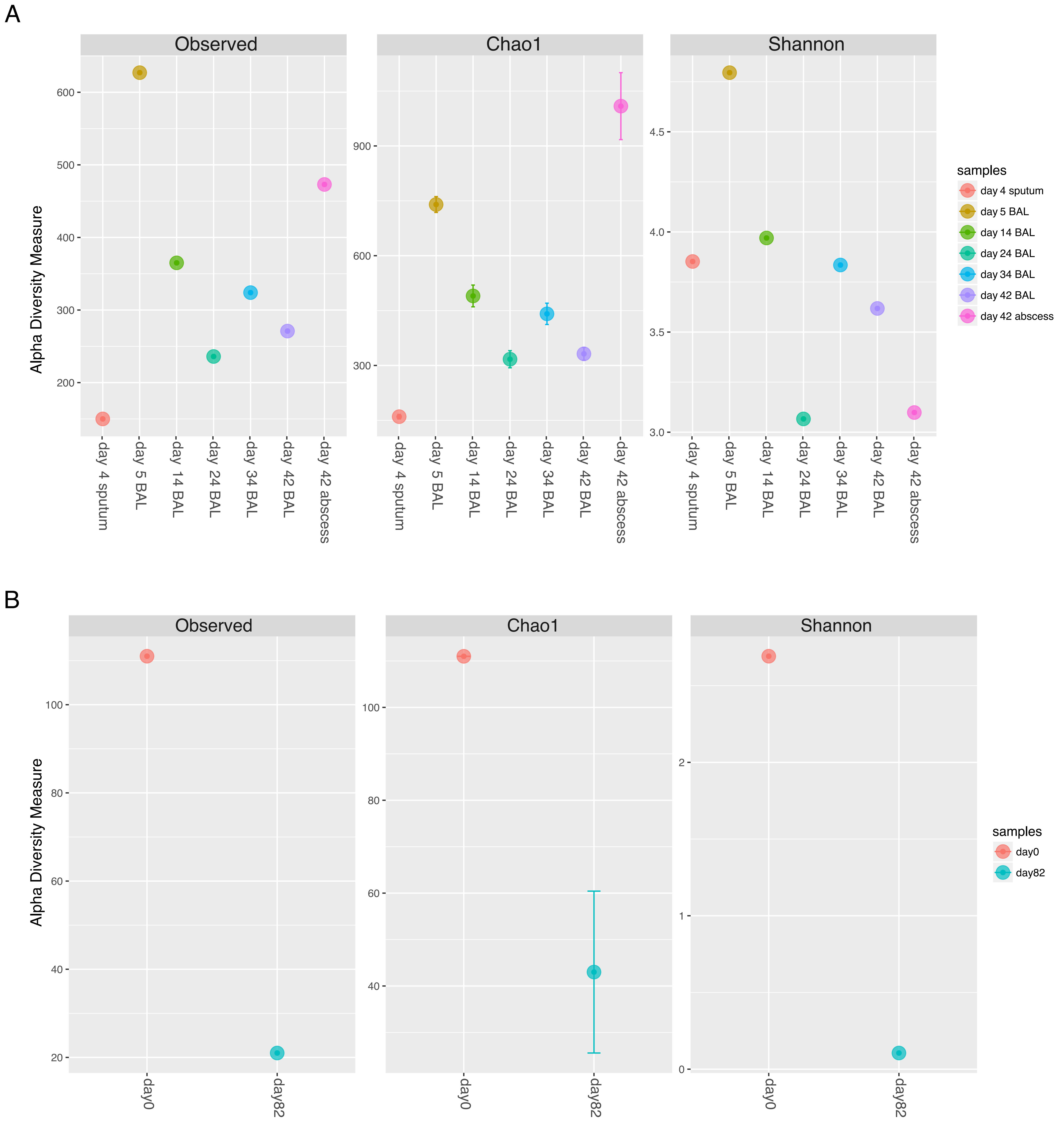
A
Observed
Chao1
Shannon
Alpha Diversity Measure
600
500
400
300
200
900
600
300
4.5
4.0
3.5
3.0
day 4 sputum
day 5 BAL
day 14 BAL
day 24 BAL
day 34 BAL
day 42 BAL
day 42 abscess
samples
B
100
80
60
40
20
2
1
0
day0
day82

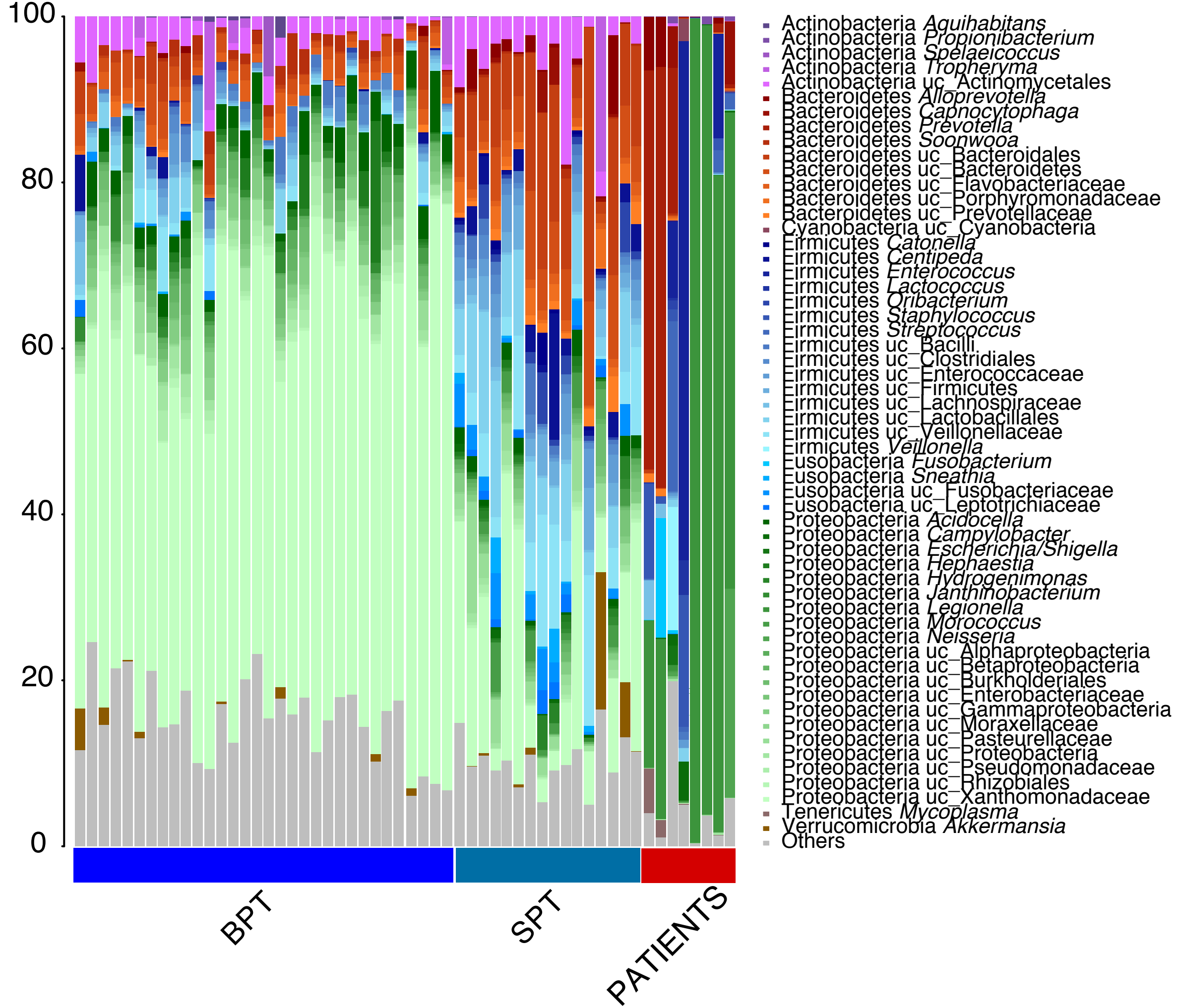

100
80
60
40
20
0
Actinobacteria Aquihabitans
Actinobacteria Propionibacterium
Actinobacteria Spelaeicoccus
Actinobacteria Tropheryma
Actinobacteria uc_Actinomycetales
Bacteroidetes Alloprevotella
Bacteroidetes Capnocytophaga
Bacteroidetes Prevotella
Bacteroidetes Soonwooa
Bacteroidetes uc_Bacteroidales
Bacteroidetes uc_Bacteroidetes
Bacteroidetes uc_Flavobacteriaceae
Bacteroidetes uc_Porphyromonadaceae
Bacteroidetes uc_Prevotellaceae
Cyanobacteria uc_Cyanobacteria
Firmicutes Catonella
Firmicutes Centipeda
Firmicutes Enterococcus
Firmicutes Lactococcus
Firmicutes Oribacterium
Firmicutes Staphylococcus
Firmicutes Streptococcus
Firmicutes uc_Bacilli
Firmicutes uc_Clostridiales
Firmicutes uc_Enterococcaceae
Firmicutes uc_Firmicutes
Firmicutes uc_Lachnospiraceae
Firmicutes uc_Lactobacillales
Firmicutes uc_Veillonellaceae
Firmicutes Veillonella
Fusobacteria Fusobacterium
Fusobacteria Sneathia
Fusobacteria uc_Fusobacteriaceae
Fusobacteria uc_Leptotrichiaceae
Proteobacteria Acidocella
Proteobacteria Campylobacter
Proteobacteria Escherichia/Shigella
Proteobacteria Hephaestia
Proteobacteria Hydrogenimonas
Proteobacteria Janthinobacterium
Proteobacteria Legionella
Proteobacteria Morococcus
Proteobacteria Neisseria
Proteobacteria uc_Alphaproteobacteria
Proteobacteria uc_Betaproteobacteria
Proteobacteria uc_Burkholderiales
Proteobacteria uc_Enterobacteriaceae
Proteobacteria uc_Gammaproteobacteria
Proteobacteria uc_Moraxellaceae
Proteobacteria uc_Pasteurellaceae
Proteobacteria uc_Proteobacteria
Proteobacteria uc_Pseudomonadaceae
Proteobacteria uc_Rhizobiales
Proteobacteria uc_Xanthomonadaceae
Tenericutes Mycoplasma
Verrucomicrobia Akkermansia
Others
BPT
SPT
PATIENTS

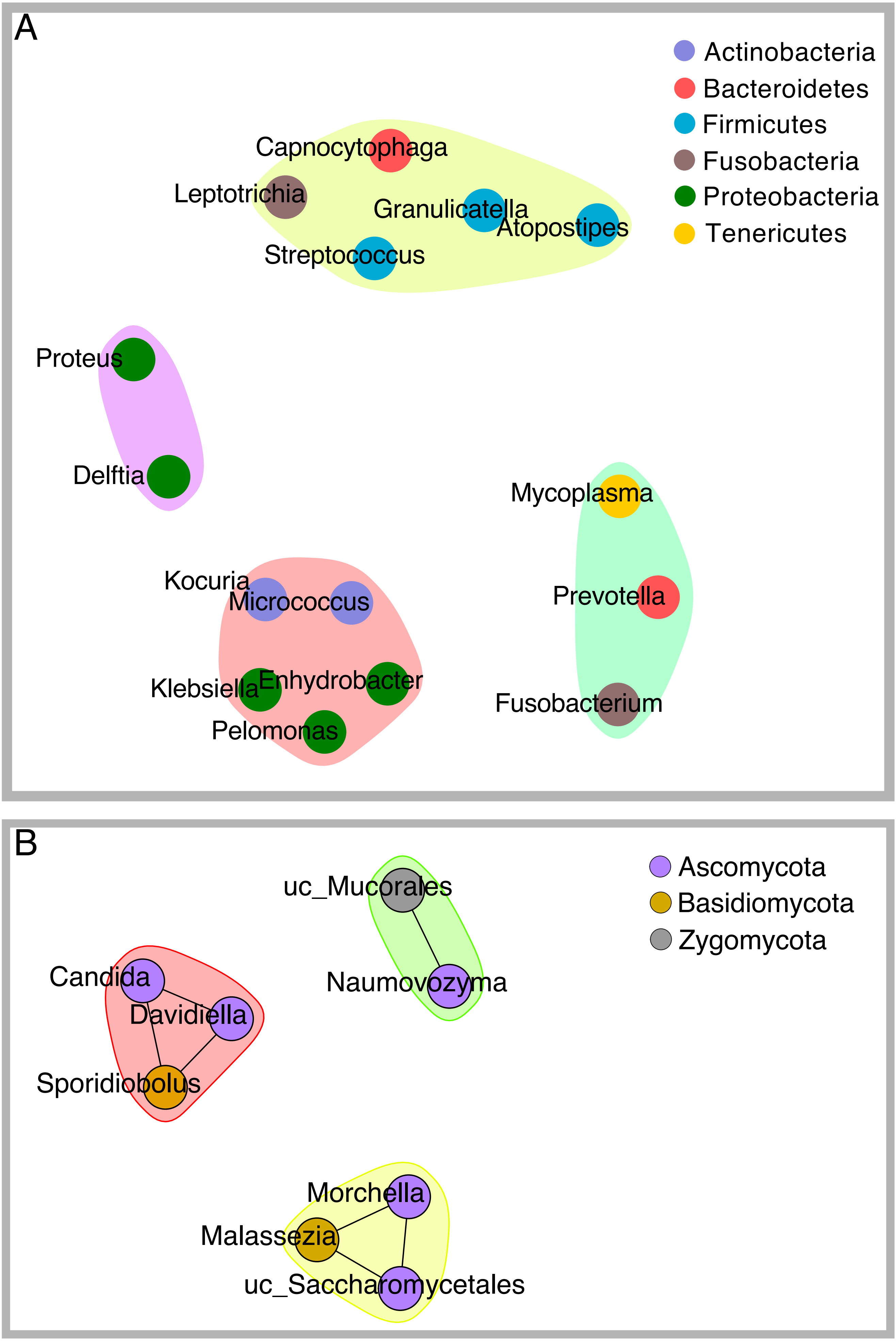
A
Actinobacteria
Bacteroidetes
Firmicutes
Fusobacteria
Proteobacteria
Tenericutes
Capnocytophaga
Leptotrichia
Granulicatella
Atopostipes
Streptococcus
Proteus
Delftia
Mycoplasma
Kocuria
Micrococcus
Prevotella
Klebsiella
Enhydrobacter
Pelomonas
Fusobacterium
B
Ascomycota
Basidiomycota
Zygomycota
uc_Mucorales
Candida
Naumovozyma
Davidiella
Sporidiobolus
Morchella
Malassezia
uc_Saccharomycetales

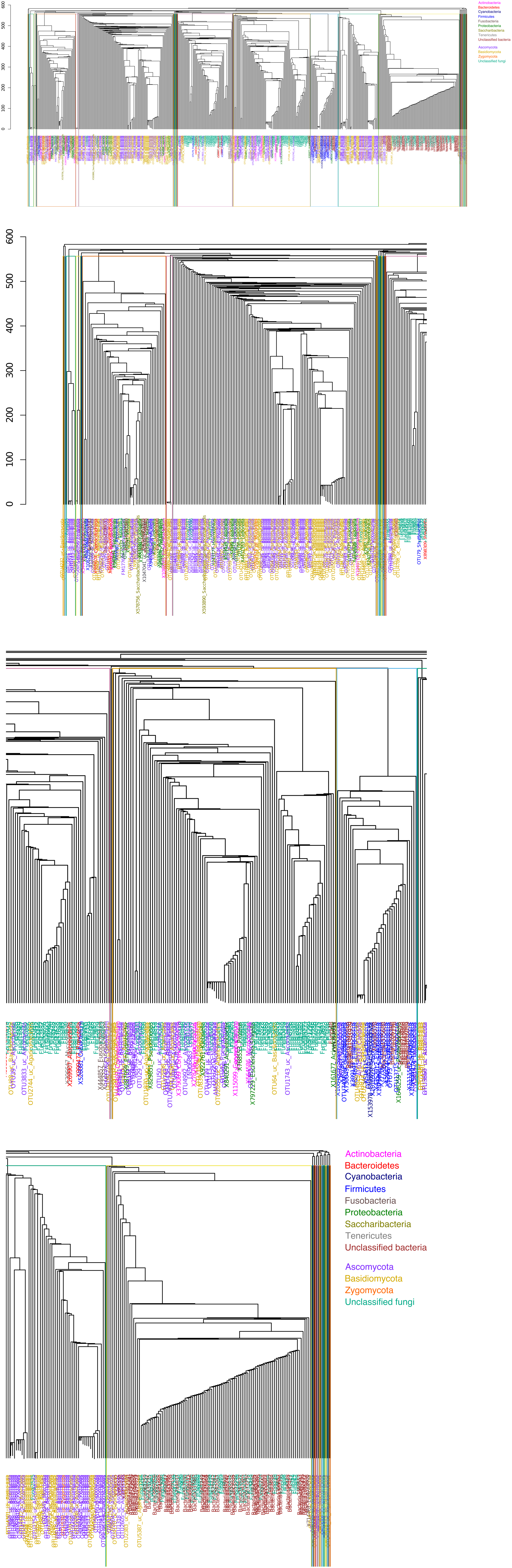

600
500
400
300
200
100
0
Actinobacteria
Bacteroidetes
Cyanobacteria
Firmicutes
Fusobacteria
Proteobacteria
Saccharibacteria
Tenericutes
Unclassified bacteria
Ascomycota
Basidiomycota
Zygomycota
Unclassified fungi
